# Long-Term Intestinal Epithelial Remodeling Induced by Acute Protein-Energy Malnutrition

**DOI:** 10.1101/2025.10.20.683425

**Authors:** Fenja A. Schuran, Neha Mishra, Víctor A. López-Agudelo, Nina Sommer, Joana P. Bernardes, Alesia Walker, Finn Hinrichsen, Ting Gong, Felix Gilbert, Lena Schröder, Archana Bhardwaj, Sven Künzel, Saskia Weber-Stiehl, Go Ito, Florian Tran, Matthieu Groussin, Christoph Röcken, Juan Matute, Stefan Schreiber, Josef M. Penninger, Richard S. Blumberg, Philippe Schmitt-Kopplin, John F. Baines, Felix Sommer, Philip Rosenstiel

## Abstract

Protein-energy malnutrition (PEM) is a global health burden with lasting effects that extend well beyond the initial nutrient deficiency. To systematically investigate the long-term effects of a single episode of PEM on the structure and function of the intestinal epithelium and its associated microbiota, we employed a comprehensive multi-omics approach, including (spatial) transcriptomics, DNA methylation analysis, fecal metagenomics, and metabolomics. Our findings show that PEM persistently alters the intestinal epithelium by depleting Paneth cells and suppressing antimicrobial gene expression - changes linked to DNA methylation that persist despite dietary recovery. In germ-free mice, the sustained epithelial phenotype after was absent. We identified the microbial lipid metabolite 9-HODE and epigenetically deregulated PPAR-driven GDF15 expression as key molecular drivers of the persistent PEM-induced Paneth cell dysfunction. Targeting microbial lipid production and its link to the host GDF15 pathway could offer novel therapeutic strategies for long-term consequences of malnutrition and other Paneth cell-associated diseases.

## Introduction

Undernutrition is a major global health burden and is responsible for 45% of child deaths world-wide ^1,2^. It is defined as a disordered nutritional state resulting from a combination of inflammation and a negative nutrient balance, leading to changes in body composition, function and outcome and associated disorders ^3^. Clinically, undernutrition has several facets, ranging from a mild caloric and/or micronutrient deficit to more severe forms resulting from sustained inadequate uptake of protein (protein energy malnutrition, termed PEM hereafter). The latter is the main form of global undernutrition and combines an energy deficit with lack of essential amino acids, which are necessary building blocks for new proteins and regulate cellular energy homeostasis. PEM is by far the most important causative factor of morbidity and mortality in developing countries and crisis regions, but also in industrialized countries the growing socioeconomic gap and specifically increasing poverty at young and old age, are linked to a significant burden of this disease state ^4^. The gut microbiota has been convincingly shown to be involved in the etiology of PEM ^5–8^ and microbiota-directed nutritional interventions have been utilized to overcome acute periods of malnutrition ^9–12^.

Whereas the pathophysiology of acute PEM has been intensely studied ^7^, long-term consequences of PEM and the molecular factors that contribute to such effects are not sufficiently understood. These consequences are characterized by a “memory effect” on specific body functions, which can be established after just one episode of malnutrition and may even be inherited by future generations^13^. Malnutrition during fetal development or early life can lead to ongoing poor weight gain and growth (known as stunting), which often persists even when a normal diet is introduced during childhood. Other clinical features associated with previous episodes of PEM include delayed motor ^14^ and intellectual development ^15–17^ and predisposition to obesity ^18^ and Type 2 diabetes in adulthood ^19–21^. Importantly, individuals who suffered from chronic malnutrition in the past are also more vulnerable to intestinal infections and have a higher mortality rate due to infection-related complications ^22,23^. Previous PEM may extend the duration of infectious diarrhea ^24^, potentially due to a decrease of intestinal barrier function, facilitating translocation of commensal bacteria and pathogens. We had previously shown that a lack of ACE2/BOAT1-dependent absorption of tryptophan in intestinal epithelial cells of mice may be a pivotal link between malnutrition, commensal bacteria and intestinal inflammation ^25^, but the duration and the exact cellular mechanism of this effect remained unclear. Early studies have shown that PEM may sustainably hinder the growth of epithelial cells in the small intestinal crypts, causing delayed cellular migration along the crypt-villus axis ^26^. This may result in a delay in the restoration of the epithelial structure and the recovery of the enzymatic and absorptive capacity in malnourished individuals ^27^. Moreover, serum anti-antibodies to intestinal microbial antigens (e.g. flagellin, LPS) are persistently elevated after PEM, indicating an insufficient barrier function and sustained immune responses against the gut microbiota ^28–30^. Despite this wealth of observations pointing towards fundamental long-term effects of PEM on the intestinal mucosa, the effector mechanisms, e.g. microbe-derived metabolites acting on cell fate decisions, remains largely unexplained. Here, we employed a murine model of adult PEM to systematically understand long-term effects of severe episode of protein undernutrition on the structure and functionality of the intestinal mucosa and its associated microbiota.

## Results

### Persistent consequences of experimental protein energy malnutrition in the small intestine of adult mice

Protein-restricted isocaloric diets in mice have been suggested to resemble the acute phenotype of protein energy malnutrition ^23,31,33^. To establish a dietary PEM model in mice, we first conducted a pilot study to assess the ability of isocaloric experimental diets with different protein levels (0.7%, 2%, 5% protein, hereafter termed “PEM diet”) to cause a PEM phenotype, as determined by the maximum weight loss within a four-week treatment period, compared to a standard diet with 20% protein, which is termed “control or ctrl diet” hereafter (Fig S1A). To investigate the long-term physiological effects of an episode of protein-energy malnutrition (PEM) on the intestinal tract, we conducted a dietary study using 10-week-old adult male C57 BL/6J mice. The mice were first subjected to a 4-week isocaloric PEM diet containing 0.7% protein, followed by a 6-week recovery phase on a regular control diet with 20% protein. (Figure 1A). After 4 weeks of 0.7% protein diet, mice lost approx. 25% of body weight compared to their littermates fed with control diet. After switching the recovery group to the control diet following 4 weeks of PEM, the mice experienced rapid weight gain during the first 2 weeks. However, by the end of the 6-week follow-up period, they had not fully regained the weight levels of the mice that had been maintained on the control diet for the entire duration of the study. Notably, this weight difference emerged after the mice had completed their axial growth, suggesting a lasting, “memory-like” effect of the prior PEM period in adult mice (Figure 1B). In a separate experiment, we confirmed that the dietary regimen successfully induced stunting in 4-week-old mice. However, we opted to proceed with the adult model to avoid potential secondary effects of perturbed developmental processes (Figure S2). No significant difference in food consumption was detected between the study groups (Figure 1C). As expected, PEM induced transient effects in several organ systems. We noticed decreased liver, epididymal white adipose tissue (EWAT) and spleen weight after 4 weeks of PEM diet (Figure S1B), which were however completely restored to the levels of their littermate controls after the shift to the control diet. Strikingly, exposure to PEM also led to several persistent changes in the intestinal tract. We observed a significantly reduced length of the small intestine in acute PEM and after recovery. While only subtle changes of crypt-villous architecture (crypt depth and villous length) were detectable (Figure S1C), persistently reduced Ki-67 and TUNEL-positive epithelial cells indicated a slower proliferation and turnover of the small intestinal epithelial lining as sequelae of PEM as shown previously in humans ^26,27^. This effect was limited to the small intestine, the primary site for protein digestion and absorption. In contrast, the colon showed reduced length (Figure 1D) and impaired epithelial proliferation (data not shown) only during the acute phase of PEM, both of which fully normalized after 6 weeks on the control diet.

**Figure 1:**
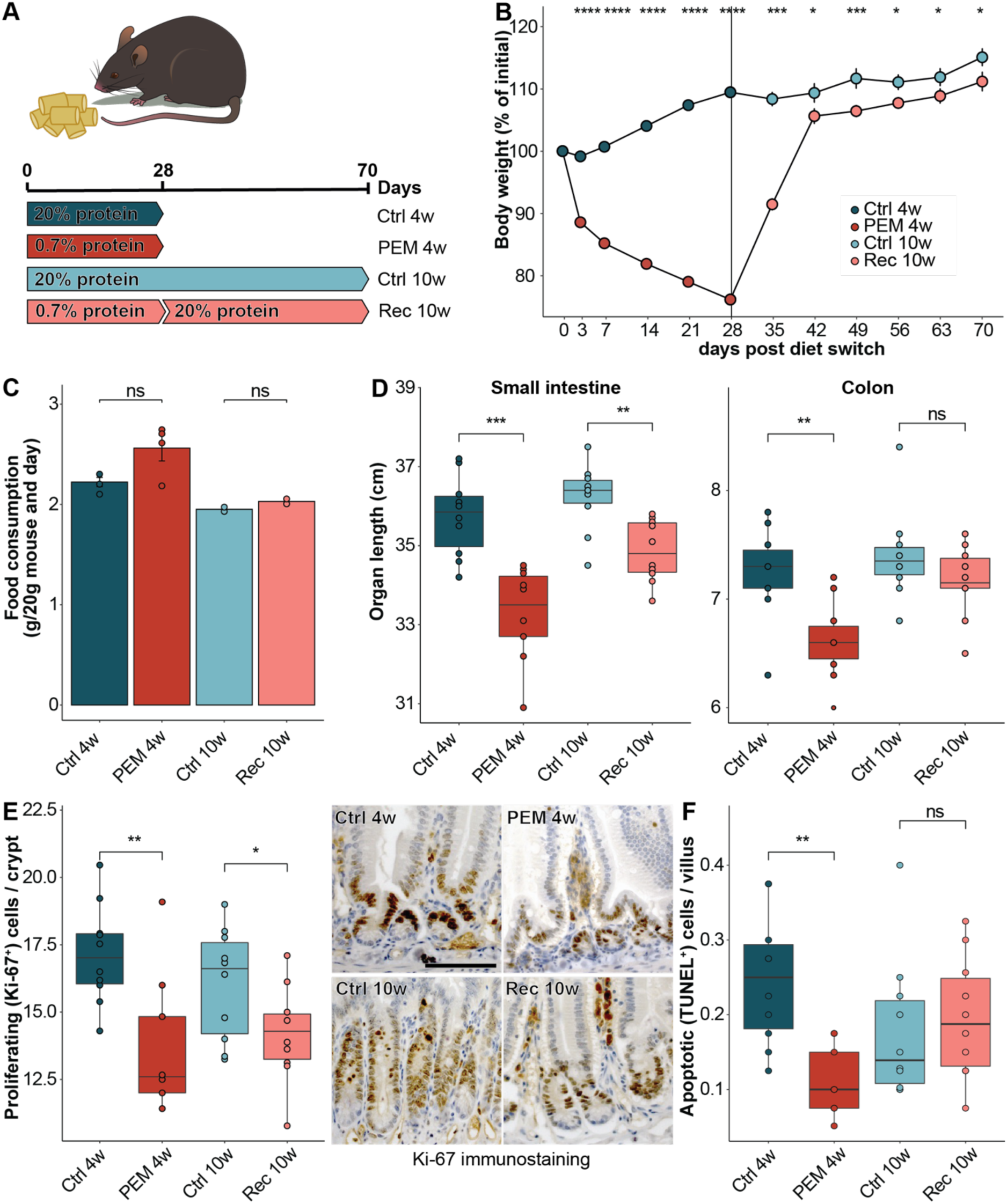
PEM induces long-lasting changes in intestinal physiology independent of caloric intake. **A)** Scheme of the PEM regimen in mice (n=10 mice per group). 12-week-old C57BL6 mice were given either isocaloric control or protein-reduced food for 28 days to induce an acute PEM, and the PEM-mice were then switched to control diet as well for an additional 42 days to monitor recovery from PEM. **B)** Body weight curve shows drastic weight loss during PEM and incomplete recovery after switch to control diet. *p < 0.05, ***p < 0.001 and ****p < 0.0001 using two-way ANOVA. **C)** Food intake of the experimental groups. ns = not significant using Mann-Whitney U test. **D)** Lengths of the small and large intestine during PEM and recovery. PEM shortened both the small and large intestine and the small intestinal length could not be restored during recovery. **p < 0.01, ***p < 0.001 and ns = not significant using Mann-Whitney U test. **E-F)** Fewer proliferating cells per crypt **(E)** and apoptotic cells per villus **(F)** in PEM mice as determined by Ki-67 immunostaining and TUNEL assay including representative images. The scale bar represents 50 µm. *p < 0.05, **p < 0.01 and ns = not significant using Mann-Whitney U test.

### Acute PEM induces a complex transcriptional dysregulation and leads to sustained downregulation of anti-microbial defense modules in intestinal epithelial cells

We next aimed to delineate the dynamics of the transcriptional response of intestinal epithelial cells during PEM and after recovery. We performed RNA sequencing of purified intestinal epithelial cells from the small intestine at acute PEM (4 weeks) and after 6 weeks recovery (10 weeks). Principal component (PC) analysis showed a strong separation between the acute PEM samples and all other conditions on the first PC (Figure S3A) indicating a severe alteration of global transcriptome signatures of the epithelium under acute PEM. A total of 8,197 genes were differentially expressed as a result of PEM, with 4,190 downregulated and 4,007 upregulated, either during the acute phase or after recovery, compared to controls. 4,125 of these differentially expressed genes (DEGs) were unique to the acute PEM phase only (2,007 down- and 2,118 upregulated genes). The upregulated PEM-specific DEGs included genes involved in key metabolic processes, such as acyl-CoA dehydrogenases (Acadl, Acadm, Acadvl, Acads, Acad11), phosphokinases (Khk, Tkfc), and glutathione transferases (Gsta1, Gsta2, Gsta3, Gsta4, Gstm1, Gstm2, Gstm3, Gstm4, Gstm6, Gstp1, Gstp2, etc.). Gene set enrichment analyses identified an over-representation of metabolic processes, including beta-oxidation, glyceraldehyde-3-phosphate metabolism, and glutathione homeostasis, likely reflecting a significant metabolic shift and heightened oxidative stress in response to PEM. (Figures 2A, S3D). Among the downregulated DEGs, changes comprised decreased innate immune and TNF production, such as *Cd14*, *Cd2*, *Ccr2*, *Ccr5*, *Ccr7*, *Il17a*, *Stat3*, *Tlr2* among others (Figures 2A, S3D).

**Figure 2:**
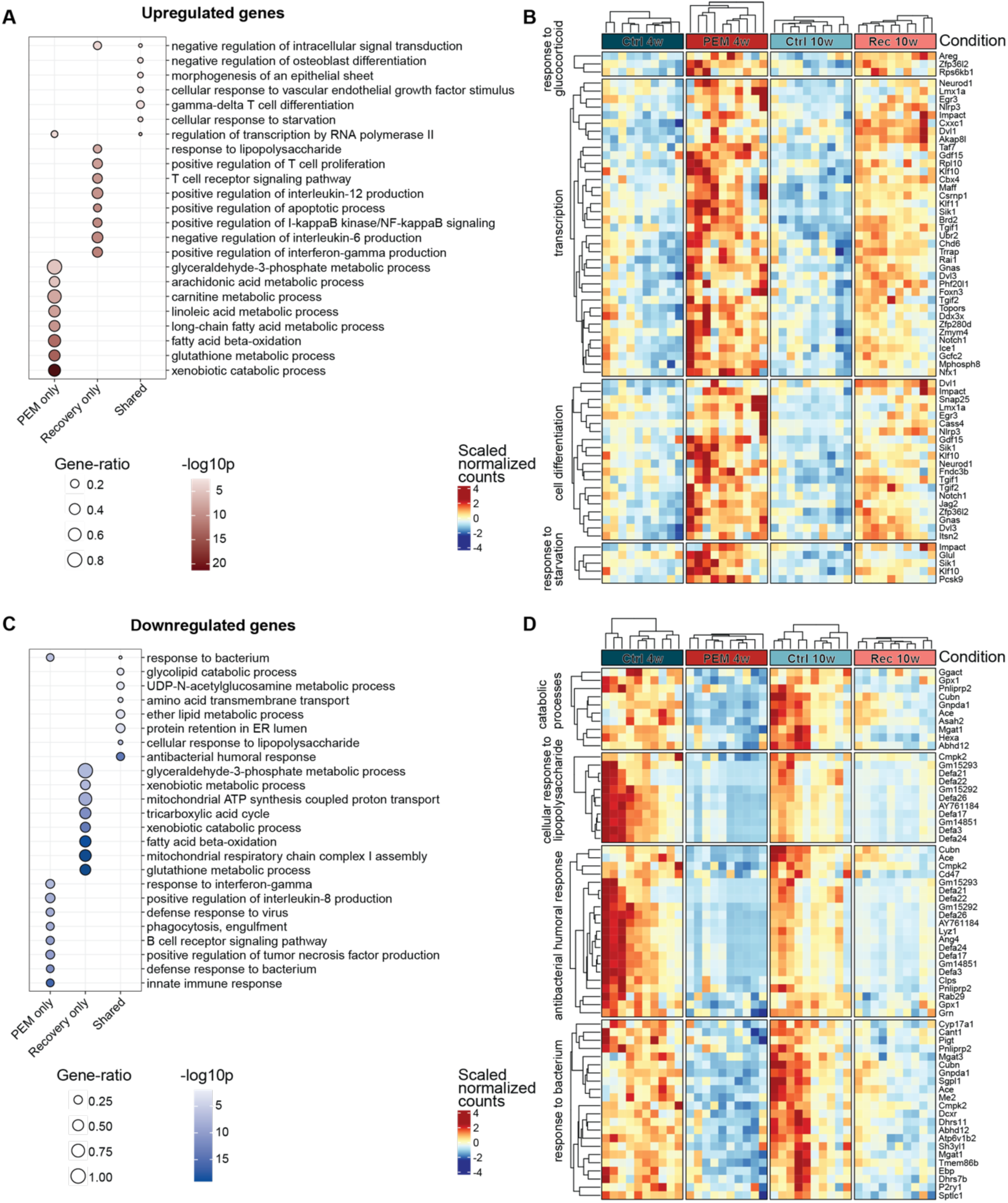
Transcriptome analysis of purified intestinal epithelial cells reveals lasting antimicrobial and metabolic alterations. **A, C.** Gene ontology (GO) terms enriched in upregulated (**A**) and downregulated (**C**) genes only at 4 weeks (PEM only), 10 weeks (Recovery only) and at both 4 and 10 weeks (Shared). Dot size is proportional to the gene ratio and color corresponds to the p-value of enrichment. Top selected terms are visualized. **B, D.** Heatmaps showing the expression of DEGs corresponding to selected GO terms enriched in upregulated (**B**) and downregulated (**D**) shared genes. Scaled normalized gene expression counts across all samples are plotted. Note that due to the GO setup, genes can belong to multiple GO categories.

Following recovery, 3,791 genes were uniquely differentially expressed in post-PEM animals compared to age-matched controls, with 2,011 genes downregulated and 1,781 upregulated. (Figure S3B). Several DEGs followed an inverse pattern at acute PEM (4w) and recovery (10w) suggesting compensatory regulatory processes after recovery from a period of protein restriction, e.g. downregulation of fatty acid beta-oxidation and glutathione metabolism-related DEGs relative to age-matched controls in gene set enrichment analyses (Figure S3D).

A comparison of the two timepoints to identify potential transcriptional “PEM-memory” effects revealed an overlap of 281 DEGs (172 downregulated and 109 upregulated; Figure 2 and S3B). These genes exhibited consistent transcriptional regulation in the same direction during both the acute phase of PEM (4 weeks) and after recovery (10 weeks), compared to age-matched controls. Upregulated shared DEGs were enriched for differentiation-related transcripts including *Dvl1* and *Dvl3* (Dishevelled Segment Polarity Protein 1 and 3), *Notch1* (Neurogenic locus notch homolog protein 1), *Neurod1* (Neuronal Differentiation 1), which pointed towards sustained alteration of epithelial lineage decisions (Figure 2B). Persistently downregulated DEGs comprised a large body of antimicrobial effector molecules such as several alpha-defensins (*Defa3, 17, 21, 22, 24, 26*), epithelial lysozyme (*Lyz1*) and intelectin (*Itln1*), indicating an ongoing reduction of epithelial defense mechanisms after PEM (Figure 2D). To further explore the topographical distribution of epithelial alterations, we performed spatial transcriptomics on intestinal tissues isolated from mice under acute PEM (4 weeks) and age-matched controls (Figure 3C). Spots containing greater than 60% epithelial tissue within the capture zone were manually selected and classified according to their anatomical position, namely “Crypt”, “Villus-Intermediate” (including both bottom and middle sections) and “Villus-Tip”, using high-resolution images of H&E-stained intestinal Swiss roles (Figure 3C, S3F). When comparing these regions between control and PEM mice, we found that the upregulated metabolic processes in PEM were primarily localized to the villi (Figure 3D), whereas downregulation of antimicrobial effector molecules (*Defa3, 5, 17, 23, 26*) was confined to the crypt region (Figure 3E).

**Figure 3:**
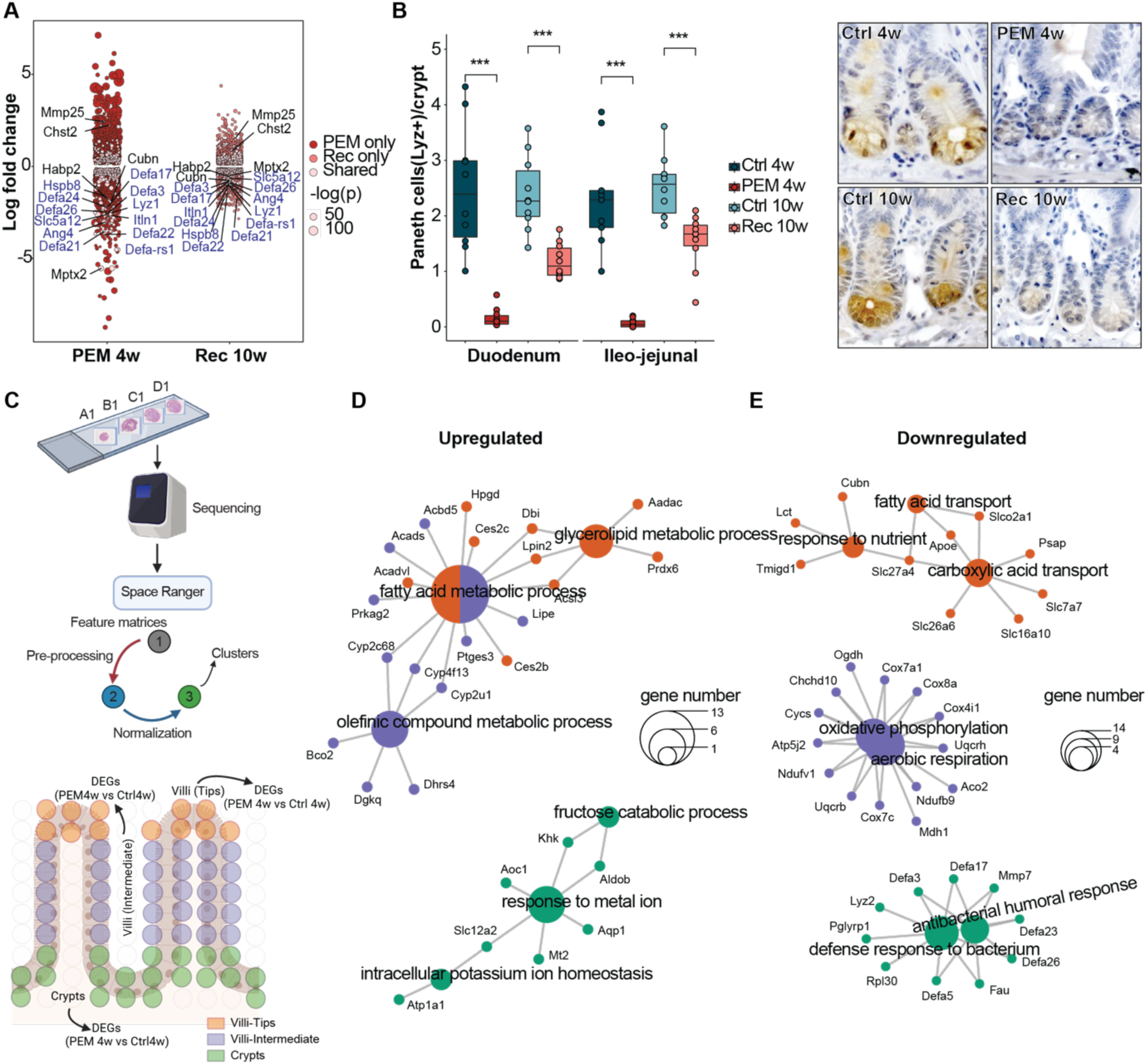
PEM impairs Paneth cell differentiation and leads to spatially confined gene expression changes along the crypt-villus axis. **A)** Log2-fold change (y axis) of differentially expressed genes (DEGs) between mice under Ctrl and PEM diet at 4 weeks and 10 weeks (x axis) with n=10 per group. Color discriminates genes that are upregulated or downregulated only at 4 weeks (PEM only), 10 weeks (Recovery only) and at both 4 and 10 weeks (Shared), and point size represents statistical significance (adjusted p value). Paneth cell related genes are highlighted in blue. **B)** Boxplots showing average number of paneth cells (Lyz+) per crypt in duodenum and ileo-jejunal regions of the small intestine of mice under Ctrl and PEM diets along with representative histological images. ***: *p* < 0.001 Ctrl_4w versus PEM_4w and Ctrl_10w versus Rec_10w using Mann-Whitney U test. **C)** Classification of spots into multiple regions: crypts represented in green, intermediate villi in purple, and villi tips in orange. Identified spots were used for DEGs analysis across PEM vs Ctrl conditions. **D-E)** Top enriched gene ontology terms of significant upregulated **(D)** and downregulated **(E)** DEGs discovered in crypts are presented. DEGs were sorted into “up” (log2foldchange > 0) and “down” (log2foldchange < 0) categories with an adjusted p-value (padj) < 0.05. Similar steps of gene ontology analysis were then applied to the Villi-Intermediate and Villi-Tip regions.

As many of the genes involved in the transcriptional memory effect of PEM, such as Defa3, Defa17, Defa21, Defa22, Defa24, Defa26, as well as Ang4, Hspb8, Itln1, and Lyz1 (Figure 2D, 3A), are recognized as Paneth cell marker genes, we next enumerated small intestinal Paneth cells by lysozyme immunohistochemistry. Indeed, numbers of lysozyme-positive Paneth cells were significantly reduced in the acute phase of PEM and remained at a much lower level after recovery (Figure 3B). Proportions of Goblet cells, another cell type of the secretory lineage, were unaffected (Figure S3C).

To test the long-term stability of PEM-dependent gene signatures, we isolated intestinal organoids from mice during the PEM-recovery diet experiment at acute PEM (4 weeks) and after recovery (10 weeks) including from CTRL diet-fed mice. These organoids were cultured for at least five passages in regular organoid medium (no nutritional restriction), before RNA was isolated and subjected to RNA sequencing. Comparing the general expression patterns between intestinal epithelial cells isolated directly from the mice and cultured organoids revealed a largely similar expression pattern both under acute PEM (Figure 4A) and after recovery (Figure 4B). The persistence of these changes in the *ex vivo* setting suggests that a single episode of PEM induced fixed regulatory alterations, leading to the observed long-lasting transcriptional changes.

**Figure 4:**
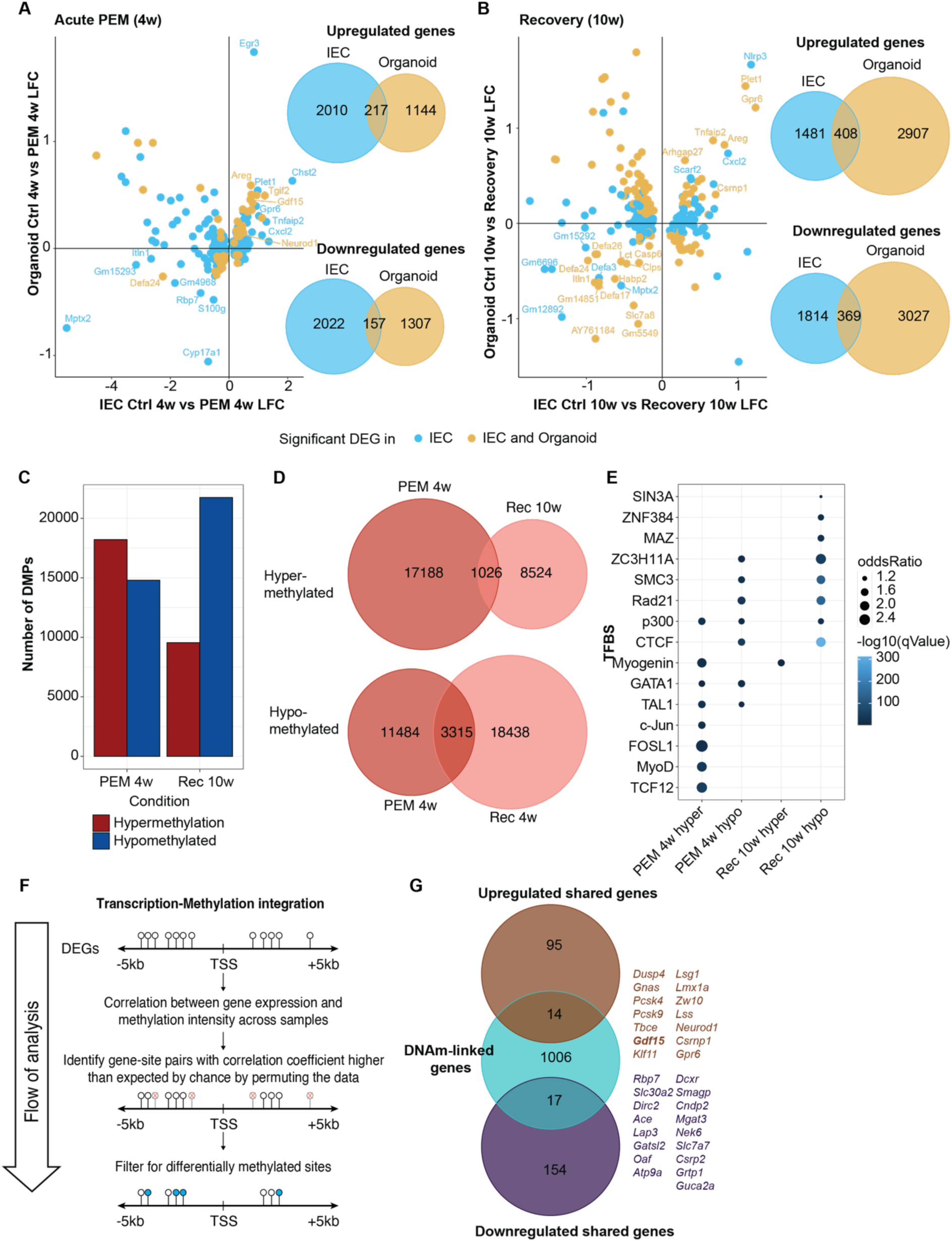
PEM-dependent persistent DNA methylation changes and *ex vivo* stability of gene expression changes. **A-B)** Comparison of transcriptional PEM effects in intestinal epithelial cells (IEC, blue) and organoids (yellow) isolated of mice from two separate dietary PEM-recovery intervention experiments during **(A)** acute PEM and **(B)** after recovery. The dot plot depicts the Log_2_[fold-changes PEM_4w/10w vs Ctrl_4w/10w] whereas the Venn diagram depicts the overlap in up- and downregulated DEGs. **C)** Number of differentially (hypo- and hyper-) methylated positions (DMPs) detected by BeadCHiP array in small intestinal epithelial cells from mice during acute PEM (4 weeks) and after recovery (10 weeks) and respective controls. **D)** Shared DMPs between acute PEM (4 weeks) and recovery (10 weeks). **E)** Predicted transcription factor binding sites enriched in observed DMPs. Dot size is proportional to the odds ratio and color corresponds to the p-value of enrichment. Top selected transcription factors are visualized. **F)** Schematic depiction of gene expression - DNA methylation integration analysis. Spearman’s rank correlation coefficient between the methylation intensity and gene expression values was calculated between the DMPs located within a range of 5kb before or after the transcription start site of the respective DEG. **G)** Venn Diagram depicting the overlaps between shared DEGs between PEM_4w and Rec_10w and DEGs correlated with their nearby DMPs, identifying 31 candidate DEGs linked with DMPs including GDF15 that has been linked to body mass regulation.

### Integrative Analysis of Genome-Wide DNA Methylation and Differential Gene Expression in Small Intestinal Epithelial Cells During Acute PEM and Post-Recovery

DNA methylation is an epigenetic mechanism that plays a key role in the long-term regulation of gene expression. In our previous work, we demonstrated its involvement in the differentiation processes of intestinal epithelial cells ^32,34^. To find a potential mechanism of the observed long-lasting transcriptional memory effect after PEM, we next analyzed genome-wide DNA methylation signatures by bead-based arrays using purified intestinal epithelial cells from the small intestine at acute PEM (4 weeks) and after recovery (10 weeks) and respective controls. In the acute PEM samples, we identified 33,013 differentially methylated positions (DMPs), of which 18,214 were hyper- and 14,799 hypomethylated (Figure 4C). Whereas after recovery, hypomethylation was the prevailing pattern compared to age-matched controls (21,753 hypo- vs. 9550 hypermethylated sites). Similar to the intersection of transcriptome signatures shared by acute PEM and recovery samples, we found that a notable proportion of DMPs from acute PEM overlapped with DMPs detected in the recovery group, with 1026 hyper- and 1007 hypomethylated sites being persistently deregulated (Figure 4D). Differentially methylated sites significantly overlapped with binding motifs for several transcription factors, for example the cell division proteins and PPAR targets RAD21 and SMC3 ^35,36^, as well as the mRNA transporter ZC3H11A and transcriptional repressor CTCF which were enriched in hypomethylated sites shared between acute PEM and recovery samples (Figure 4E).

To delineate regulatory DNA methylation changes linked with persistently altered gene expression *in cis*, we integrated transcriptome and methylome data using a hierarchical approach as described ^32,37^ (Figure 4F). First, we located DMPs situated within a range of 5kb before or after the transcription start site of any DEG. Next, we computed the correlation between the expression level of each DEG and the intensity of methylation at the corresponding DMPs. Among 56,271 deregulated DMPs analyzed, 6,554 were located within the defined window of at least one DEG. Out of these, 6,197 DMP-DEG pairs representing 3,394 genes exhibited significant correlation, with 51% displaying a reciprocal correlation (i.e., high methylation linked to low transcript levels or vice versa) and 49% exhibiting a cooperative correlation (i.e., high methylation linked to high transcript levels or vice versa). 31 DMP-DEG with a persistent pattern in acute PEM and after recovery were identified (Figure 4G). We were able to identify persistent links between DNA methylation and transcriptome signatures, both after acute PEM and in the recovery phase. The cis-linked DNA methylation-DEG pairs point towards potential long-term alteration of cellular states through epigenetic regulation after a PEM episode. Notably, GDF15 (Growth/Differentiation Factor 15) emerged as a promising candidate for mediating some of the long-term PEM memory effects, given its established role as an endocrine signal of nutritional stress and body mass regulation and as a mediator of cancer cachexia ^38–40^.

### Intestinal microbial community dynamics in acute PEM and after recovery indicate a high level of resilience to PEM-induced perturbation

To explore the potential involvement of gut microbial dysbiosis for the observed sustained phenotypes, we performed 16S rRNA gene sequencing on sequential stool samples collected from mice during the PEM-recovery experiment (Figure 5A). During PEM (up to week 4), bacterial α-diversity as measured by the Shannon Index declined significantly compared to control-fed mice, but fully normalized again after switching to control diet during recovery (from week 5 to 10) (Figure 5B). Composition of the fecal microbiota from mice of under acute PEM differed profoundly from baseline (before dietary intervention) or control-fed littermates (both at 4 weeks and 10 weeks ”), and PEM-recovery mice at 10 weeks (PERMANOVA on between-sample Bray-Curtis dissimilarities, F-statistics = 25.5, R^2^ = 0.54, p = 1e-4, Figure 5C). Specifically, in mice subjected to PEM, *Enterobacteriaceae*, a family of the class Proteobacteria that includes opportunistic pathogenic microorganisms such as *Escherichia* and *Shigella*, which have been linked with infection-related pathologies in malnourished individuals ^41,42^, were expanded along with *Bacteroides*, *Parabacteroides* or *Lachnospiraceae* UCG-006. In contrast, the relative abundance of *Bifidobacterium*, *Bilophila*, *Lachnospiraceae* NK4A136, *Lachnoclostridium*, *Alistipes*, *Parasutterella*, *Faecalibaculum* and *Muribaculaceae* were significantly decreased (Wilcoxon rank sum test, p < 0.001) compared to control-fed mice (Figures 5D and S4A, S4B). No significant differences were observed at the taxonomical level between PEM-recovery mice and their littermate controls. Notably, the intraclass correlation coefficient (ICC), which measures the similarity among samples within the same group, was significantly higher in fecal samples from mice with acute PEM compared to all other experimental groups (Figure S4D). This suggests that acute PEM induces uniform changes in the microbiota, with minimal influence from individual baseline condition or other environmental factors. Through shotgun metagenomic sequencing at weeks 4 and 10, we also explored changes of the functional potential of the microbiome. At 4 weeks, we found that a wide range of metabolic functions related to fatty acid and amino acid metabolism were enriched in the microbiome of mice under acute PEM compared to control-fed mice. (Figure 5E and S4C). Again, after recovery at week 10, no significant changes compared to control mice were observed, suggesting that adult microbial communities exhibit strong resilience to this type of perturbation.

**Figure 5:**
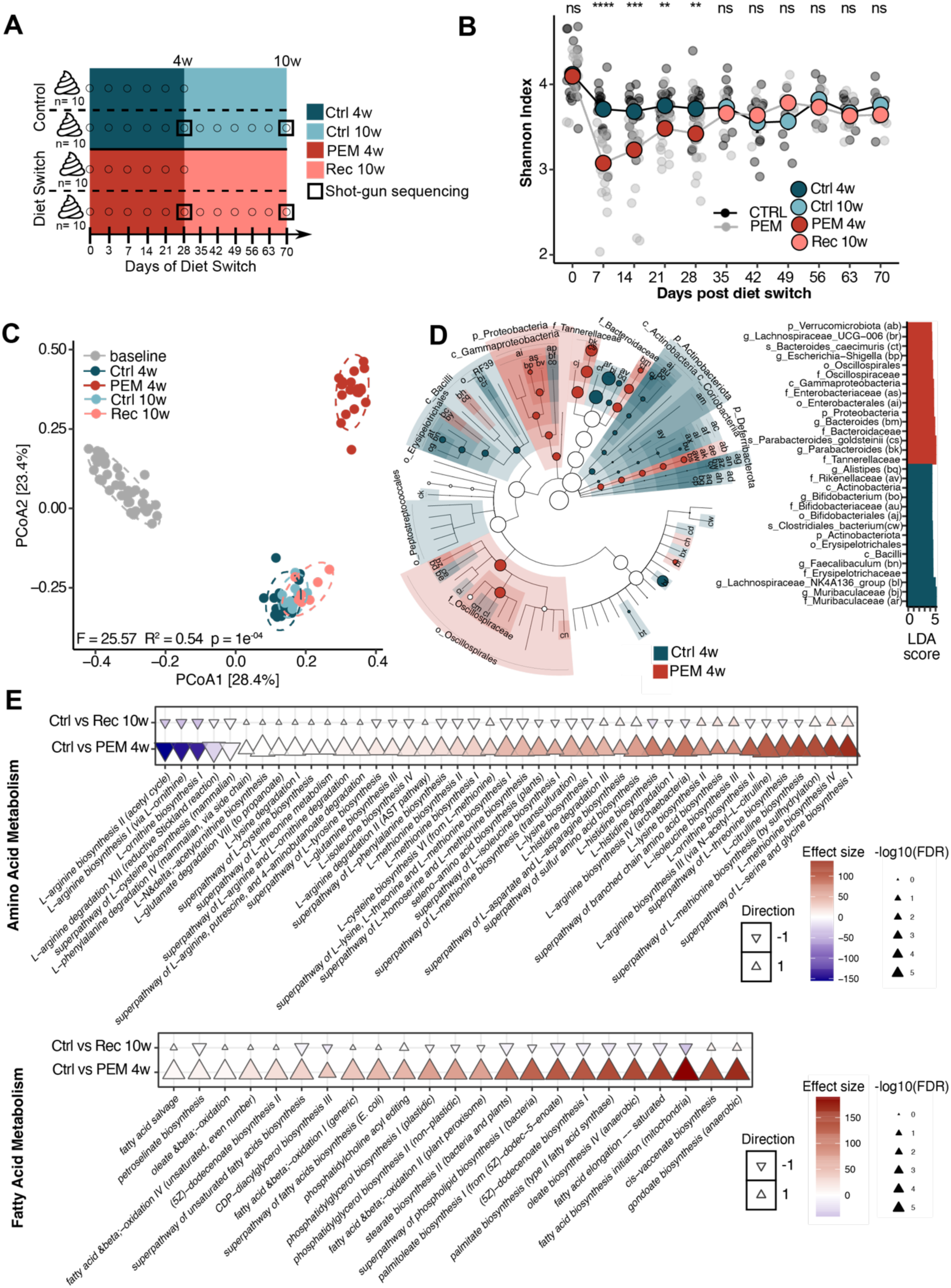
Acute PEM drastically alters microbiome composition and function. **A)** Stool sample collection scheme for microbiome analyses. Stool samples were collected weekly during the 10 weeks of diet experiment (normal or reduced protein diet) and used for 16S amplicon sequencing analysis and taxonomy characterization. Shot-gun metagenomic sequencing and functional potential analysis was performed only on stool samples from weeks 4 and 10 (Ctrl_4w, PEM_4w, Ctrl_10w and Rec_10w, each n=10 per group). **B)** Longitudinal alpha diversity of stool samples from mice under normal and reduced protein chow. Shannon index was computed at ASV level. Longitudinal cross-sectional comparisons were performed using Wilcoxon signed rank test. ns: not significant, *: *p* < 0.05, **: *p* < 0.01, ***: *p* < 0.001, ****: *p* < 0.0001. **C)** Principal Coordinate Analysis based on Bray-Curtis’s distances of stool fecal microbiomes of mice during PEM and recovery to explore further differences in microbiome composition. Differences in the centroids of groups were tested with PERMANOVA with 10000 permutations. **D)** Linear discriminant analysis (LDA) effect size (Lefse) of the fecal microbiomes at peak acute PEM (day 28, PEM_4w versus Ctrl_4w). The cladogram depicts the phylogenetic distribution of differential taxa and the bar plot depicts differential taxa ranked by LDA. Differences are shown by the color of the most abundant class, with red indicating PEM_4w and blue indicating Ctrl_4w. The diameter of each circle corresponds to the abundance of the taxon. This analysis uses the SILVA v138 taxonomy. The bar plot represents the top 30 taxa based on LDA scores. The letters in brackets denote the abbreviation of the taxon shown in the cladogram. **E)** Differential amino acid and fatty acid metabolic microbiome pathways (PEM_4w vs Ctrl_4w and Rec_10w vs Ctrl_10w). The triangles show the hues and direction of the effect size (b coefficient). Color intensity and size represent the magnitude of the effect size and FDR significance of the specific pathway, respectively.

### Lack of an intestinal microbiome ameliorates phenotype severity and alters fecal metabolite patterns upon PEM perturbation

We next explored the role of the microbiome in shaping PEM-related phenotypes in our mouse model. We hypothesized that, similar to other stressors like DSS or radiation-induced enteritis ^43^, the absence of intestinal microbes—which induces an energy-restricted state, particularly in the intestinal mucosa—would exacerbate PEM symptoms. We thus performed the same PEM-recovery diet intervention in germfree (GF) mice of the same genetic background. Contrary to our expectation, GF mice exhibited significantly less body weight loss than conventionally raised (CONV-R) mice with a normal microbiome (Figure 6A). PEM-induced body weight loss could be fully normalized during recovery after switching to a control diet in GF, but not in CONV-R mice (Figure 6A). Importantly, in contrast to CONV-R mice, numbers of Paneth cells were not affected by PEM in the absence of a microbiota (Figure 6B). Organ measures and abundance of other cell types were also unaltered by PEM in GF mice (Figure S5). Untargeted metabolomics of fecal samples from GF and CONV-R mice during acute PEM (day 28, week 4) revealed that lipid metabolites were significantly impacted by the nutritional challenge in a microbiota-dependent manner (Figure 6C). Specifically, conjugated linoleic acids and oxylipins such as 12,13-diHOME and 9-HODE were strongly induced under acute PEM in CONV-R mice, but were virtually absent in GF mice as individually validated by targeted quantitative metabolomics (Figure 6C, D). Supporting this metabolomic data, abundances of the respective functional genes in the microbiome, namely of lipoxygenases (LOX) and epoxide hydroxylases (EH), which generate 9-HODE and 12,13-diHOME from linoleic acid, respectively, were increased under acute PEM only in CONV-R mice as revealed by shotgun metagenomics (Figure 6E). Importantly, using publicly available metagenomic data of malnourished children undergoing a nutritional intervention with either a Microbiota Directed Complementary Food (MDCF-2) or standard Ready-to-Use Supplementary Food (RUSF) ^10^, we found that abundances of microbial LOX and EH genes declined during the therapeutic nutritional intervention (Figure 6F). These findings collectively demonstrate that fecal oxylipins, such as 9-HODE and 12,13-diHOME, along with the microbial enzymes that produce them, are highly sensitive to protein-energy malnutrition (PEM).

**Figure 6:**
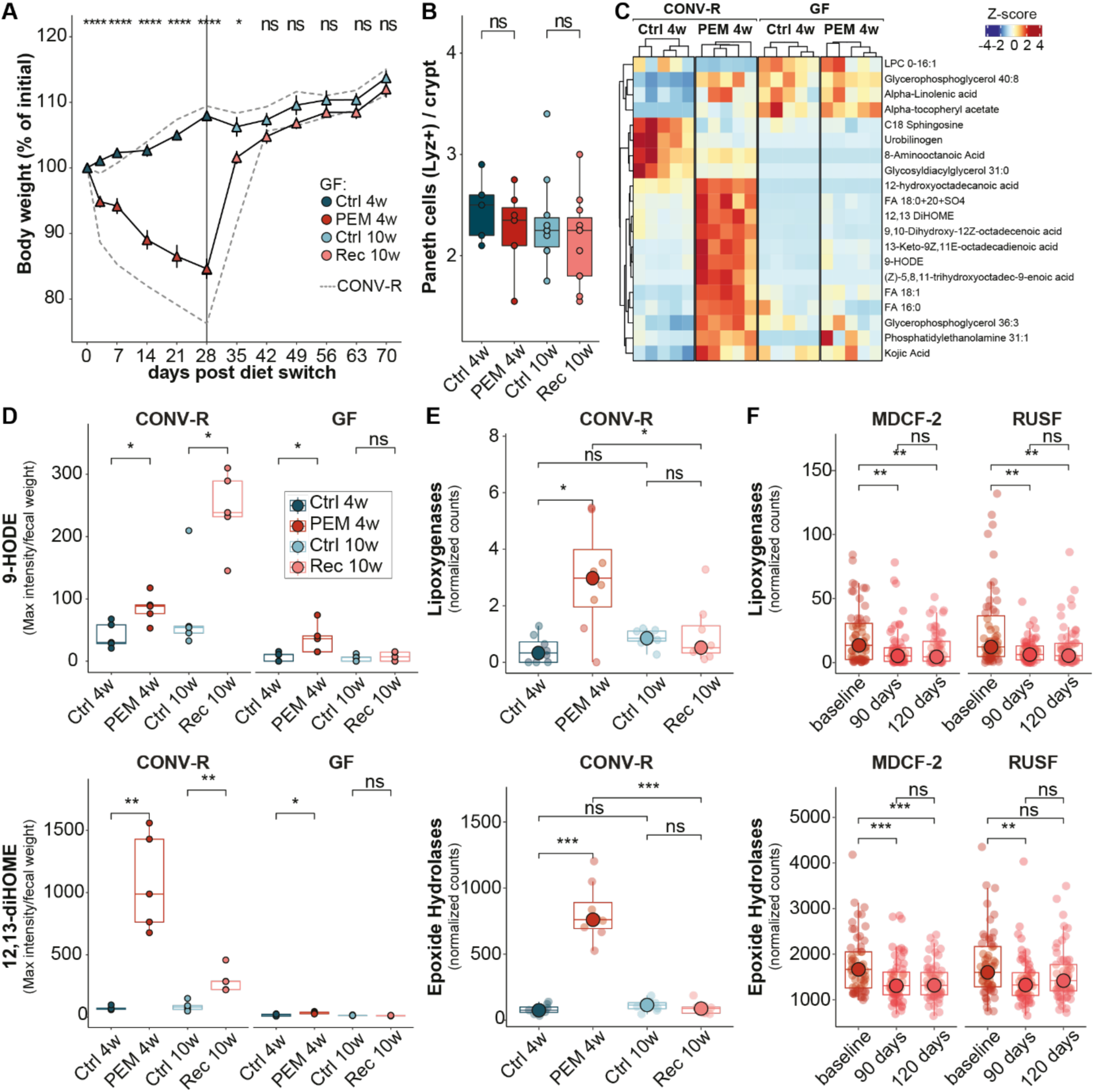
Absence of microbial signals attenuates the Paneth-cell phenotype and fecal metabolic alterations upon PEM. **A)** Body weight curves of GF mice without a microbiome under 4-week PEM regimen followed by 6-week recovery with regular diet compared to CONV-R mice with a regular microbiome. Note that GF mice displayed milder disease symptoms with significantly less body weight loss during PEM and a full recovery (n=10-11 mice per group). **B)** Paneth (LYZ1-positive) cell count in small intestinal sections of GF mice. **C)** Top 20 most significant lipid metabolites identified in feces of GF and CONV-R mice fed a CTRL or PEM for 28 days. **D)** Levels of 9-HODE and 12,13diHOME in feces of GF and CONV-R mice during the PEM & recovery experiment. **E)** Bacterial lipoxygenase and epoxide hydrolase (EH) genes identified in stool metagenomics of GF and CONV-R mice during PEM & recovery. **F)** Lipoxygenase and EH genes in feces collected on day 0, 90 and 120 of malnourished children undergoing a nutritional intervention with either a Microbiota Directed Complementary Food (MDCF-2) or standard Ready-to-Use Supplementary Food (RUSF) using published metagenomics data ^10^. Data in (A) were compared by two-way ANOVA and in (B) and (D-F) using Wilcoxon signed rank test. ns: not significant, *: *p* < 0.05, **: *p* < 0.01, ***: *p* < 0.001, ****: *p* < 0.0001.

### Activation of the PPAR-GDF15 pathway disrupts Paneth cell development during PEM

Oxylipins like 12,13-diHOME and 9-HODE are recognized as peroxisome proliferator-activated receptor (PPAR) agonists, and Paneth cell differentiation is regulated by PPAR signaling. We hypothesized that these microbially-derived metabolites may serve as a mechanistic link between the gut microbiota and disruptions in Paneth cell function ^44–47^. PPARs are nuclear lipid-activable receptors controlling a variety of genes regulating metabolism, including GDF15 ^48,49^. Strikingly, we observed that the expression patterns of both *Gdf15* and *Pparg* were co-dependent on PEM and the microbiome, showing upregulation in response to PEM and during recovery in CONVR mice, but not in GF mice (Figure 7A). We next assessed the functional role of PPAR activation and GDF15 for Paneth cell differentiation. Paneth cell differentiation was induced by a switch to standard ^50^ ENR-CD medium containing CHIR99021 (a GSK3 inhibitor/WNT activator) and DAPT (a γ-secretase inhibitor) (Figure 7B). Successful induction was reflected by increased expression of Paneth cell markers over time, such as Lyz1 (Figures 7C, D, E) and Defa5 (Figure 7C). We first tested the effects of GDF15 on Paneth cells *in vitro*. Strikingly, stimulation of intestinal organoids with recombinant and TGFβ-free GDF15 (rGDF15) ^51^ resulted in downregulation of Paneth cell markers (Figure 7C). To further explore the link between PPAR signaling and GDF15, we established an *in vitro* organoid PEM model. Intestinal organoids were cultured in either normal or protein-reduced ENR medium (see methods), while simultaneously inducing Paneth cell differentiation by CHIR99021 and DAPT addition. Of note, when organoid cultures were treated with Lanifibranor, a pan-PPAR agonist, under PEM conditions, Paneth cell marker expression was significantly repressed (Figure 7D), while Lanifibranor in the absence of protein restriction had no discernible effect. Additionally, Lanifibranor increased *Gdf15* expression, confirming it as a direct downstream target of PPAR under PEM (Figure 7D). Similar effects were observed with the microbial PPAR agonist 9-HODE *in vitro* under PEM conditions. After 96 hours, 9-HODE decreased *Lyz1* expression and upregulated *Gdf15* after 48 hours (Figure 7E). The timing of events, with an initial upregulation of *Gdf15* expression followed 48 hours later by a decrease in *Lyz1* marker expression in response to 9-HODE stimulation, collectively still supports a mechanistic link. The interaction of PEM and PPAR-GDF15 activation was required to induce defective Paneth cell differentiation as stimulation with Lanifibranor or 9-HODE in the absence of PEM did not reduce *Lyz1* or *Gdf15* expression (Figure S6). These findings therefore suggest the presence of a host-intrinsic PPAR-GDF15 pathway in the intestinal epithelium which can be activated by microbially-derived oxylipin metabolites during PEM and mediates sustained effects on Paneth cell differentiation and function.

**Figure 7:**
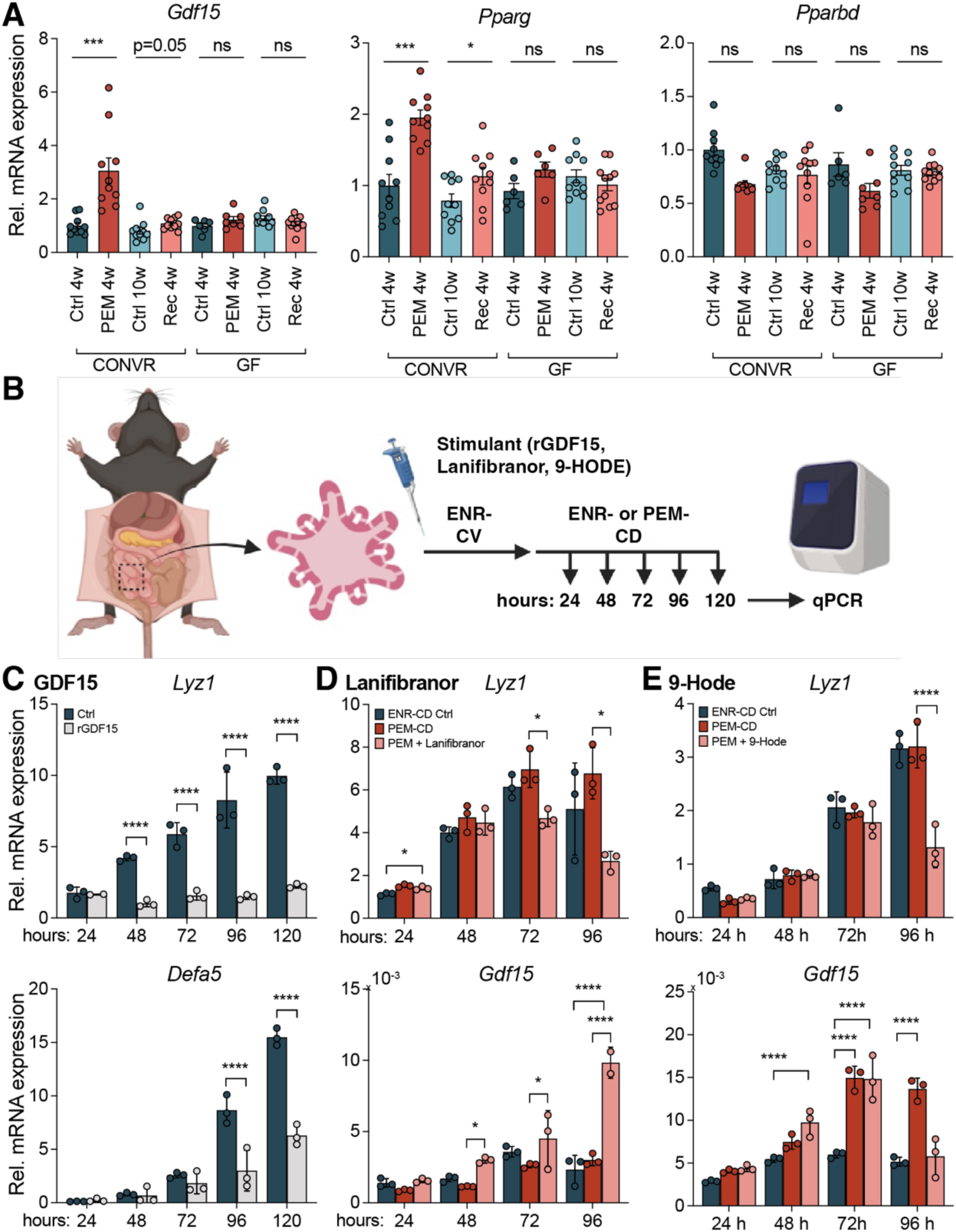
Microbiome and PEM dependent factors block Paneth cell differentiation. **A)** Expression of *Gdf15*, *Pparg* and *Pparbd* were assessed in small intestinal tissue of both CONVR and GF mice subjected to PEM and after recovery. *p < 0.05, ***p < 0.001 and ****p < 0.0001, ns = non-significant. Significance testing was performed using Wilcoxon-Mann-Whitney-Test. **B)** Schematic drawing of organoid intervention experiment. Murine intestinal organoids were first cultured in ENR-CV stem cell organoid medium and then Paneth cell differentiation was induced using ENR-CD or PEM-CD medium. Stimulants (recombinant GDF15 [1 µg/ml], pan-PPAR-agonist Lanifibranor [20 µM] or the lipid 9-HODE [1 µM] were added during initial ENR-CD and PEM-CD culture and kept throughout the experimental duration. All experiments were performed independently at least twice in triplicates. **C-E)** Relative *Lyz1*, *Defa5* and *Gdf15* mRNA expression in organoids during Paneth cell differentiation and stimulated with **(C)** recombinant GDF15, **(D)** the Pan-PPAR-agonist Lanifibranor or the lipid **(E)** 9-HODE. Note that PPAR-activation boosts GDF15 levels thereby blocking Paneth cell differentiation and the bacterial PEM metabolite 9-HODE also suppresses Paneth cell differentiation. *: *p* < 0.05, **: *p* < 0.01, ***: *p* < 0.001, ****: *p* < 0.0001 and ns = not significant using Mann-Whitney U test.

## Discussion

PEM remains a major global health challenge, impacting millions of individuals worldwide. In recent decades most studies have focused on understanding acute protein undernutrition. Major achievements are based on using locally available ingredients and co-targeting the intestinal microbiome ^5,8,10,11,25,52–57^. However, the exact nature of the interactions between the intestinal microbiota and host tissues during nutritional stress are still largely unknown. In this study, we utilized a murine model of adult PEM to systematically investigate the long-term effects of a single severe episode of protein deficiency on both the structure and function of the intestinal mucosa, as well as its associated microbiota. Using multi-omics profiling, we uncovered extensive transcriptional, epigenetic, metabolic, and cellular effects of acute malnutrition. Our findings provide a resource of mucosa-microbiota alterations in the acute phase of PEM, which significantly extends previous insights and can be leveraged to mine for novel targeted interventions ^5,8,9,11,58–61^. Importantly, we show that a single episode of PEM leads to a prolonged reduction in body weight and induces significant structural changes in the intestinal epithelium. These include macroscopic alterations, such as a shortened small intestine, and microscopic changes, such as a depletion of Paneth cells, which is a common feature of environmental enteric dysfunction (EED) in malnourished children ^62,63^. Paneth cells are key regulators of intestinal physiology and antimicrobial responses, thus, our findings of permanently impaired Paneth cell function due to PEM could potentially explain the increased susceptibility towards gastrointestinal pathogens, i.e. *Enterobacteriaceae*, accounting for severe diarrhea, transmigration peritonitis and sepsis observed in severely malnourished humans ^22,23^. The structural changes were accompanied by “memory-like” transcriptional alterations in the intestinal epithelium, notably a prolonged reduction in the expression of key Paneth cell genes, including those encoding antibacterial factors such as defensins and lysozyme, during and after acute PEM. These gene expression changes were sustainably present even in intestinal organoids grown from intestinal stem cells of mice harvested during acute PEM or recovery, when cultured for multiple passages *in vitro* under full-nutrient conditions. This suggests the involvement of long-term regulatory mechanisms, such as alterations in DNA methylation ^64,65^, which control the lasting effects of PEM. Indeed, we could show a strong remodeling of DNA methylation patterns upon PEM in purified intestinal epithelial cells. We conducted an interaction analysis between gene expression and differentially methylated positions ^32,37^ and identified, besides other differentiation factors (e.g. Neurod1 ^66^), a link between cis-linked methylation changes and upregulation of mRNA levels of the pleiotropic stress-induced cytokine GDF15 as a potential mediator of PEM on intestinal function and metabolic processes ^67–69^.

To assess the dynamics of gut microbial community shifts during and after PEM we conducted 16S rDNA and metagenomic shotgun sequencing. In line with previous studies ^5,7–9,11,58,61^ we detected a significant shift of gut microbiota composition and functional potential during acute PEM. Strikingly, PEM-dependent changes completely reverted back to their original configuration during recovery indicating a high level of resilience of fecal microbial communities^70^. We initially hypothesized that lack of microbiome would increase PEM susceptibility, e.g. due to an energetic restriction in the absence of bacterial fermentation ^71,72^ or lack of TLR-dependent protective signals^43^. Surprisingly, when subjecting GF mice to PEM, we found a significantly milder gastrointestinal phenotype with a less pronounced bodyweight loss, and neither a long-lasting reduction of bodyweight nor significant depletion of Paneth cells. These findings indicate that a second microbially-dependent hit is required to trigger a strong PEM phenotype. Using metabolomics in CONV-R and GF mice, we identified the fecal oxylipin metabolites 12,13-diHOME and 9-HODE as potential sources of the PEM- and microbiome-dependent phenotype. By analyzing metagenomic data, we found increased abundances of LOC and EH genes encoding enzymes that generate 9-HODE and 12,13-diHOME from linoleic acid under acute PEM in CONV-R mice and in malnourished children, which suggests that the same mechanism functions in mice and humans. Both, 9-HODE and 12,13-diHOME, have been implicated as immunomodulatory lipids, which may act through PPARs ^73–75^. Using an *ex vivo* malnutrition model in intestinal organoids, we then found that protein deprivation alone did not impair Paneth cell differentiation, but a second (microbial) hit targeting the PPAR/GDF15-axis was required.

Our data indicate that the sustained loss of Paneth cells could be linked to an epigenetically fixed upregulation of GDF15 in intestinal epithelial cells. Whereas the intestinal epithelium has been shown to be the major site of inducible GDF15 ^76^ upon nutritional stressors, there are conflicting results on the expression distribution of its cognate receptor glial cell line-derived neurotrophic factor family receptor α-like (GFRAL) as well as on biological effects of yet unidentified non-GFRAL GDF15 receptors. Although the specific receptor system involved was not identified in our study, we used a recombinant GDF15 source, which ruled out the possibility of co-purified TGF being responsible. This suggests that the observed effects are directly attributable to GDF15 itself. Our findings are of particular clinical relevance as biologic treatments targeting GDF15 have not only proven effective in mice ^77^, but they are currently entering phase 2 clinical trials for tumor cachexia in patients ^40,78^. While the main effect of this new drug class is to increase bodyweight via enhanced appetite, muscle function and physical performance, these drugs could also prove effective in restoring Paneth cell numbers and function in malnourished patients, thereby alleviating gastrointestinal symptoms and reducing PEM lethality. A major obstacle in a global use of biologicals for PEM will be the expected high cost of therapy. Instead, targeting the microbial metabolites 12,13-diHOME or 9-HODE could provide a cost-effective alternative significantly extending the scope of successful nutritional interventions targeting the intestinal microbiome ^10^.

### Potential limitations of the study

This study was conducted using a single episode of PEM in an adult murine model. Even though there is evidence that these findings can be translated to the human situation (e.g. Paneth cell disruption, elevated GDF15 levels, elevated abundance of LOX/EH genes), our comprehensive molecular profiling results pointing towards a long-term memory-like effect of PEM remain to be validated in a human setting. Moreover, studies on the relevance of our findings for the childhood situation, e.g. for developmental phenotypes such as stunting ^2^, are warranted. In addition, like most controlled PEM studies we used an isocaloric set-up, which does not reflect the usual undernutrition scenario in crisis regions, where both available protein and overall calories are severely reduced. It must be emphasized that the alteration of lipid metabolism, which we see both in the mucosal transcriptome and bacterial metagenome, is not linked to increased lipid content of the synthetic diet, as protein calories are replaced by carbohydrates. Since lipid metabolism has been shown to be altered in human malnutrition cohorts through both microbiome and serum analyses ^79–82^, we are confident that the observed changes are primarily driven by metabolic shifts resulting from limited protein availability. Importantly, due to our experimental setup, we do neither address the effects of multiple PEM periods nor the duration of the observed memory phenotype in the intestinal epithelium. From earlier regeneration kinetics, it seems however unlikely that the effect is simply caused by the physiological lag time of differentiation, as acute chemical ablation of Paneth cells was followed by re-appearance of mature cells already after 24h ^83^. An interesting explanation for the disappearance of Paneth cells could be a permanent trans-differentiation to stem-like dividing cells, which could partially replace the original stem cells ^84,85^. Further follow up studies including longer observation periods and/or multiple PEM episodes and single cell-based analyses are required to comprehensively study the effects on epithelial lineage decisions, but also long-term effects in other cell types and tissue compartments.

It is increasingly evident that phenotypic responses to PEM vary between the sexes ^86^. In our study, most analyses have been done in males. In the GF condition, we included both sexes and observed no significant differences between them. Further research should therefore investigate sex-dependent effects during PEM, especially since woman during pregnancy and breast-feeding are a very vulnerable group of patients prone to suffering from PEM due to an increased protein demand ^87^. Moreover, in this study we utilized a PEM model characterized by a general protein deficiency, rather than a shortage of specific amino acids or their combinations. Certain amino acids, such as tryptophan, undergo extensive metabolism by both the host and the microbiota, generating metabolites with diverse signaling roles in immune regulation and differentiation processes. These metabolites are known to influence the expression of antimicrobial peptides ^25,88^. It will be crucial to investigate whether the acute and memory-like effects of PEM can be reversed through supplementation of these amino acids or their metabolites, such as those involved in the kynurenine/nicotinamide pathway.

## Conclusions

In this study we present a comprehensive multi-omics resource of a dietary PEM model in mice, revealing that a single episode of PEM induces long-lasting phenotypic changes, with a particular focus on the intestinal mucosa and microbiota. Key findings include sustained weight loss, shortening of the small intestine, impaired proliferation of intestinal epithelial cells, a reduction in Paneth cells, and diminished expression of genes encoding antimicrobial agents. Notably, germ-free (GF) mice exhibited alleviated acute PEM symptoms and did not manifest any of the aforementioned long-term PEM-associated phenotypic changes. Through an integrative multi-omics analysis and *ex vivo* intestinal organoid stimulations, we identified microbial lipid metabolites, such as 12,13-diHOME and 9-HODE, along with the host PPAR-GDF15 axis, as potential molecular drivers of the enduring PEM phenotype, particularly the dysfunction of Paneth cells. These findings suggest that targeting microbial lipid production or the host GDF15 pathway could provide novel therapeutic strategies, not only for malnutrition, but also for other conditions associated with Paneth cell dysfunction, such as Crohn’s disease ^89,90^.

## Methods

### Animals

All animal experiments were approved by the local animal safety review board of the federal ministry of Schleswig Holstein and conducted according to national and international laws and policies (V242-30405/2016(55-5/16), V242-6900/2019(28-3/19) and V242-60285/2019(105-9/19)). C57Bl6/J mice were purchased from Charles River Laboratories and housed in the Central Animal Facility (ZTH) of the University Hospital Schleswig Holstein (UKSH, Kiel, Germany) under specific-pathogen free (SPF) conditions in individually ventilated cages (Green Line, Techniplast) or under sterile conditions in gnotobiotic flexible film isolators at the Max Planck Institute for Evolutionary Biology in Plön. All mice were kept under a 12-h light cycle and fed a regular gamma-irradiated chow diet *ad libitum*.

### Dietary PEM intervention experiment

All custom PEM and CTRL diets were purchased from Ssniff (Table S1). To induce PEM in adult mice, 10-week-old CONV-R or 10-16-week-old GF C57Bl6/J mice (n=40) were either given a control diet (CTRL, 20% protein) or an isocaloric diet with reduced protein content (PEM, 0.7% protein) for 4 weeks to induce an acute PEM. To induce PEM in young mice, 4-week-old C57Bl6/J mice (n=40) were either given a control diet (CTRL, 20% protein) or an isocaloric diet with reduced protein content (PEM, 2% protein due to a greater PEM susceptibility of young mice) for 4 weeks to induce an acute PEM. One half of each experimental group was killed for blood & organ harvest at 4 weeks to assess the acute PEM phenotype. The other half of the experimental groups was either kept on or switched to a diet with regular protein content for an additional six weeks to assess recovery from PEM. Up to five recipient mice were co-housed. Mice were weighed every week and fecal pellets were collected, immediately frozen on dry ice and stored at –80°C. For final analyses, blood was collected from the vena cava, mice were killed by cervical dislocation and tissues were removed for histological and molecular analyses.

### Histology and immunostainings

Tissue specimen were fixed in methacarn (60% dry methanol, 30% dry chloroform, 10% glacial acetic acid) for 1-2 weeks at room temperature prior to paraffin embedding and sectioning. 5 µm thick sections were cut and deparaffinized in Roti-Histol and rehydrated in ethanol followed by hematoxylin and eosin (H&E) or periodic acid-Schiff (PAS, TH Geyer, 3952016) staining. Alternatively, sections were boiled in citrate buffer for antigen retrieval and subjected to immunostaining. For this purpose, we used the Vectastain ABC kit (Vector Laboratories), peroxidase-substrate solution from DAB Peroxidase (HRP) Substrate Kit (Vector Laboratories) and primary antibodies for detecting proliferating cells were incubated in a 1:500 diluted mouse anti-Ki67 antibody (BD Biosciences, cat.no. 556003) and for Paneth cells in a 1:500 diluted mouse anti-LYZ1 antibody (Santa Cruz, cat.no. sc-27956) overnight at 4°C. TUNEL assay was performed with the *in situ* cell death detection Kit, TMR red (Roche, 12156792910) according to the manufacturer’s instructions, while nuclei were counterstained with DAPI (4’,6-diamidino-2-phenylindole, dihydrochloride) nuclear dye (Thermo Fisher, 62247). Slides were visualized using a Zeiss Imager Z1 microscope (Zeiss) and pictures were taken using ZEN pro (Zeiss) software.

### Isolation of intestinal epithelial cells (IECs)

IECs were isolated from small intestinal tissue using the Lamina Propria Dissociation Kit (Miltenyi BioTech, Bergisch Gladbach, Germany) according to the manufacturer’s protocol. In brief, intestinal epithelial cells were isolated by disruption of the structural integrity of the epithelium using ethylenediaminetetraacetic acid (EDTA) and dithiothreitol (DTT). Purity of individual IEC fractions was analyzed by flow cytometry on a FACS Calibur flow cytometer (B&D, Heidelberg, Germany) with Cellquest analysis software from Becton Dickinson. We used the Anti-EpCam-PE (Clone: G8.8, Biolegend, San Diego, USA) antibody for analysis of IEC purity.

### Noggin-conditioned medium

HEK Noggin cells, previously described by Miyoshi and Stappenbeck (2013) and provided by Prof. Zeissig’s group at CRTD, Dresden University, Germany, were used in this study. In summary, HEK cells were modified to secrete Noggin protein into the culture medium. The cells were thawed in DMEM + 10%(v/v) FBS purchased from Biochrom. Once confluent, the cells were cultured for 3 cycles in DMEM with 10% (v/v) FBS and 10 μg/mL Puromycin (Sigma-Aldrich). After the 3rd cycle, the cells were diluted 1:20 and cultured in DMEM with 10% (v/v) FBS without Puromycin. Four days later, the medium was collected and centrifuged to remove excess cells. Fresh DMEM with 10% (v/v) FBS was added and after another four days, the second batch of medium was collected and centrifuged. The medium from the first batch was then combined with the second batch.

### R-Spondin-conditioned medium

HEK R-Spondin-1-producing cells, previously described by Miyoshi and Stappenbeck (2013) and provided by Prof. Zeissiǵs group at CRTD, Dresden University, Germany, were used in this study. In summary, HEK cells were modified to secrete R-Spondin-1 protein into the culture medium. The cells were thawed in DMEM with 10% (v/v) FBS (Biochrom). Once confluent, the cells were cultured for 3 cycles using DMEM with 10% (v/v) FBS+ 300 μg/mL Zeocin (Thermo Fisher). After the 3rd cycle cells were diluted 1:20 and cultured in DMEM with 10% (v/v) FBS without Zeocin. Four days later, the medium was collected and centrifuged to remove excess cells. Fresh DMEM with 10% (v/v) FBS was added and after another four days, the second batch of medium was collected and centrifuged. The medium from the first batch was then combined with the second batch.

### Isolation and culture of intestinal organoids

Organoids were generated from small intestinal crypts of male C57Bl6/J mice receiving either a control or PEM diet following procedures described earlier by Sato et al., 2009 ^91^. Organoids were cultivated in 24 well plates at 37°C with 5% CO_2_ atmosphere in Matrigel (BD) with ENR-conditioned medium supplemented with 0.1% human recombinant EGF (50 µg/mL) as described by Sato et al., 2011 ^50^. ENR-conditioned medium consisted of 70% (*v*/*v*) 2 × basal medium (Advanced DMEM/F12 supplemented with HEPES [1M], Glutamax [100×], Penicillin/streptomycin 10,000 U/mL [1:50] and N-Acetylcysteine [500 mM]), 10% (*v*/*v*) Noggin-conditioned medium and 20% R-Spondin-conditioned medium.

### Organoid PEM model

Murine **i**ntestinal organoids were initially cultured for 48 h in ENR-CV medium, an ENR-conditioned medium containing 3.3 µM CHIR99021 (Miltenyi Biotech, #130-103-926) and 1 mM valproic acid (Sigma, PHR1061-1G) promoting differentiation into the stem cell niche. These stem cell organoids were then transitioned to either ENR-CD or PEM-CD medium containing 3.3 µM CHIR99021 and 10 µM DAPT (Tebubio, 25704-0041) to further drive differentiation into the Paneth cell niche.

The PEM culture medium (1.25% amino acids) consisted of 68.75% (*v*/*v*) Advanced DMEM/F12 (ADF) medium without amino acids or glucose [Biozol/US Biological Life Sciences, #USB-D9800-27-10] supplemented with 10 mM HEPES, 1× Glutamax, 100 U/mL Penicillin/streptomycin and 1 mM N-Acetylcysteine), 1.25% (*v*/*v*) regular ADF medium with full amino acid concentrations, 10% (*v*/*v*) dialysed Noggin-conditioned medium and 20% dialysed R-Spondin-conditioned medium. The Noggin- and 20% R-Spondin-conditioned media were dialyzed using using cellulose-hydrate tubing with a cutoff of 15 kDa (Reichelt Chemietechnik GmbH, catalogue number 84605)) to prevent the introduction of nutrients from the added conditioned media while retaining Noggin and R-Spondin required for organoid growth. for the in-vitro PEM model the conditioned Medium was dialized. Dialysis success was assessed using Multisti× 10 SG testing strips (Siemens).

### Organoid stimulations with GDF15, Lanifibranor or 9-HODE

Murine **i**ntestinal organoids were initially cultured for 48 h in ENR-CV medium, an ENR-conditioned medium containing 3.3 µM CHIR99021 (Miltenyi Biotech, #130-103-926) and 1 mM valproic acid (Sigma, PHR1061-1G) promoting differentiation into the stem cell niche. These stem cell organoids were then transitioned to either ENR-CD or PEM-CD medium containing 3.3 µM CHIR99021 and 10 µM DAPT (Tebubio, 25704-0041) to further drive differentiation into the Paneth cell niche. Organoids were divided into experimental groups and stimulated with 1 μg *E. coli* derived recombinant mouse GDF15 (R&D Systems, 8944-GD) per 500 µl organoid medium for up to 120h, 20 μM of the pan-PPAR agonist Lanifibranor (MedChemExpress, HY-104049) for 24h to 96h or 1 μM 9(S)-HODE (Sigma Aldrich, SML0503) for 24h to 96h. Corresponding control groups received vehicle treatments identical to those used for the stimuli. All organoids were then cultured as described above.

### RNA isolation and qPCR

RNA was extracted using the RNeasy Mini Kit (QIAGEN) according to the manufacturer’s protocol. Total RNA concentration was determined with a NanoDrop ND-1000 spectrophotometer (PeqLab Biotechnologie). Total RNA was reverse transcribed to cDNA using the High-Capacity cDNA Reverse Transcription Kit (ThermoFisher Scientific, 4368814). qPCR was carried out using TaqMan^®^ Gene Expression Assays (ThermoFisher Scientific) (Table S2) according to the manufacturer’s instructions on a Viia 7 Real-Time PCR System (ThermoFisher Scientific). Relative gene expression was normalized to housekeeper gene *Actb* (β-actin) and calculated by the 2^-ΔCt^ method.

### Transcriptional profiling by RNA sequencing

Total RNA from IECs and intestinal organoids was extracted using the RNeasy Mini Kit (Qiagen) according to the manufacturer’s protocol. RNA concentration and integrity were analysed using a TapeStation 4200 System (Agilent) and a Qubit 4 fluorometer (ThermoFisher Scientific). RNA-sequencing libraries were prepared using the TruSeq® RNA seq Library Prep Kit v2 according to the Illumina TruSeq® messenger (mRNA) sequencing protocol. The RNA-seq libraries were sequenced on an Illumina HiSeq 4000 sequencer (Illumina, San Diego,CA) with an average of 23 million reads (1x 50 bp) at IKMB NGS core facilities. The nf-core rnaseq pipeline (version 1.2) (https://github.com/nf-core/rnaseq) ^92^ was used to pre-process the sequencing data. Adapter and low-quality bases from the sequencing reads were trimmed using Trim Galore (0.5.0) (http://www.bioinformatics.babraham.ac.uk/projects/trim_galore/) and reads shorter than 35bp after trimming were removed. The filtered genes were mapped to the mouse genome (GRCm38) using STAR aligner (version 2.6.1a) ^93^. The expression counts of the genes were estimated using featureCounts (version 1.6.2) ^94^ and were normalized across samples using the DESeq normalization method. Differentially expressed genes (DEGs) between Ctrl_4w and PEM_4w samples as well as Ctrl_10w and Recovery 10w samples were identified using the Bioconductor package DESeq2 (version 1.20.0) ^95^. Genes with FDR adjusted p-value of 0.05 were regarded as differentially expressed (DEGs). Gene ontology enrichment analyses were conducted using the Bioconductor package topGO (version 2.46.0) ^96^ with genes with similar expression level as the universe set. In the topGO analysis, the Fisher.elim p-value, calculated using the weight algorithm, of 0.05 was used as the significance threshold.

Transcription factor binding sites (TFBS) enriched in the promoter regions of DEGs were identified by conducting enrichment analysis using the Bioconductor package LOLA (version 1.24.0) ^97^. Promoter regions were defined as the region between 1,500 bp upstream to 500 bp downstream of the transcription start site.

### Spatial transcriptomics

Small intestinal tissue was prepared for spatial transcriptomics using the Visium Spatial Fresh Frozen Protocol CG000240 (RevE, 10x Genomics). Briefly, small intestinal tissue was rolled into Swiss rolls, embedded in O.C.T. medium (Tissue-Tek) within Cryomolds (Tissue-Tek), and then rapidly immersed in pre-cooled isopentane on liquid nitrogen until the O.C.T. medium solidified. The samples were subsequently cryopreserved at −80°C. To determine the optimal permeabilization time of 15 minutes, serial 10µm sections were cut and mounted on a Visium Tissue Optimization Slide (10x Genomics). For spatial transcriptomics, additional 10µm-thick sections were mounted on a Visium Gene Expression Slide for Fresh Frozen (10x Genomics), followed by methanol fixation and H&E staining following protocol CG000160 (RevD, 10x Genomics). Whole slide imaging was conducted using a NanoZoomer S60 Digital Slide Scanner (Hamamatsu, C13210-04) with 40x magnification. Spatial library construction was carried out according to the manufacturer’s protocol CG000239 (RevH, 10x Genomics), and the libraries were then sequenced on a NovaSeq 6000 sequencer (Illumina) with 2× 100bp read length. Sequencing data was mapped to the mouse reference genome (mm10) using 10x spaceranger v.2.1. Feature-barcode matrices for each sample were imported into the R package ‘Seurat’ v.4.2.0 for quality control, normalization, dimensionality reduction, and Louvain clustering ^98^. As quality control, spots were filtered if they had <200 features (genes) or >25% of mitochondrial gene expression. After spot filtration, sample level normalization was performed using SCTransform ^99^ function in Seurat, which uses a regularized negative binomial regression to transform the UMI count data. Thereafter, sample integration was performed with the CCA algorithm followed by a Shared Nearest Neighbor (SNN) construct graph and clustering using default settings. Using high-resolution images of H&E-stained intestinal Swiss roles, multiple regions were identified, and spots classified as belonging to “Crypt”, “Muscle”, “Villus-Intermediate” (including both bottom and middle sections), or “Villus-Tip” regions. The dominant clusters corresponding to each category were then used for differential gene analysis. The “FindMarkers” function in Seurat v4.2.0 was used to identify differentially expressed genes (DEGs) within each cluster corresponding to Crypts, Villus-Tips, and Villus-Intermediate under both CT and PEM conditions. Genes were classified as upregulated if they had log2 fold changes >0 and as downregulated if they had log2 fold changes <0 with an adjusted p-value <0.05. Gene ontology enrichment analysis of these DEGs was performed using clusterProfiler v4.12.1 ^100^ with thresholds of adjusted p-value <0.05 (hypergeometric test).

### DNA methylation profiling

DNA was isolated from IECs using the DNeasy Blood and Tissue Kit (Qiagen) according to the manufacturer’s protocol. DNA methylation levels were measured using the Infinium Mouse Methylation BeadChip (Illumina), which covers > 285k methylation sites spanning over CpG islands, transcription start sites, enhancers, and gene regions. DNA methylation data was analysed using the Bioconductor package RnBeads (version 2.12.1) ^101^. Sites that overlapped with SNPs and had unreliable measurements were filtered out. Context-specific probes, probes on the sex chromosomes, and probes with missing values were also removed resulting in the removal of 54,378 out of 261,600 probes. The signal intensity values were normalized using the dasen method. Differentially methylated positions (DMPs) and regions (DMRs) between Ctrl_4w and PEM_4w samples as well as Ctrl_10w and Recovery 10w samples were identified using the automatically selected rank cutoff of RnBeads. Predicted transcription factor binding sites (TFBS) enriched in DMPs were identified by conducting enrichment analysis using the Bioconductor package LOLA (version 1.24.0) ^97^.

### Integration of DNA methylation and transcriptome data

To integrate gene expression data with DNA methylation data, we used a hierarchical approach as described before ^32,102^. We first identified DMPs located 5,000 bp upstream and downstream of the transcription start sites of DEGs. Spearman’s rank correlation coefficient between the normalized expression count of each DEG and the methylation intensity (β-values) of its corresponding DMPs were calculated. To test the statistical significance of the correlations, we calculated the false discovery rate (FDR) using a permutation approach.

### 16S rRNA gene amplicon sequencing

DNA was isolated from fecal material using the DNeasy PowerSoil Kit (Qiagen) following the manufacturer’s protocol. Extracted DNA was eluted from the spin filter silica membrane with 100 µl of elution buffer and stored at −80 °C. 16S profiling and MiSeq sequencing was performed as described earlier ^103,104^, with the following modifications: the V3-V4 region of the 16S rRNA gene was amplified using the dual barcoded primers 319F (ACT CCT ACG GGA GGC AGC AG) and 806R (GGA CTA CHV GGG TWT CTA AT) ^105^. Each primer contained additional sequences for a 12 base Golay barcode, Illumina adaptor and a linker sequence ^106^. PCR was performed using the Phusion Hot Start Flex 2X Master Mix (NEB) in a GeneAmp PCR system 9700 (Applied Biosystems) and the following program (98°C for 3 min, 25-30x (98°C for 20 s, 55°C for 30 s, 72°C for 45 s), 72°C for 10 min, hold at 4°C). Performance of the PCR reactions was checked using agarose gel electrophoresis. Normalization was performed using the SequalPrep Normalization Plate Kit (Thermo Fisher Scientific, Darmstadt, Germany) following the manufacturer’s instructions. Equal volumes of SequalPrep-normalized amplicons were pooled and sequenced on an Illumina MiSeq (2 x 300 nt). Quality control of the MiSeq 16S amplicon sequence data was performed in MacQIIME v1.9.1 (http://www.wernerlab.org/software/macqiime) ^107^. Sequencing reads were trimmed keeping only nucleotides with a Phred quality score of at least 20, then paired-end assembled and mapped onto the samples using the barcode information. Microbiome phylogenetic analyses were conducted using QIIME2 ^108^ (release 2022.8). The paired-end sequences were filtered, denoised, and chimera removed using DADA2 ^109^ plugin resulting in abundance tables of amplicon sequence variants (ASVs) with on average 7.7K ± 4.2K (mean ± sd) reads per sample. Taxonomy was assigned using the Bayesian classifier provided in DADA2 and using the Silva rRNA database v.138 ^110^. ASVs not assigned to “Bacteria” or annotated as “Mitochondria”, “Chloroplast”, or “Eukaryote” were removed. QIIME2 outputs (feature table, taxonomy assignation, and rooted tree), were combined into a single object using the qiime2R (0.99.6) and Bioconductor phyloseq R (1.40.0) ^111^ packages.

### Metagenomic sequencing

A total of 32 fecal DNA samples (8 Ctrl_4w, 8 Ctrl_10w, 8 PEM_4w, and 8 Recovery_10w), which were already included in the 16S profiling, were also subjected to shotgun metagenomic sequencing performed at the Competence Centre for Genomic Analysis (Kiel). DNA libraries were generated using the Illumina DNA Prep kit following the manufacturer’s instructions. Libraries were then pooled and sequenced on an Illumina NovaSeq 6000 with 2 x 150 nt for 19.6 ± 4.14 (mean ± sd) million reads per sample. Shot-gun reads were processed using an in-house established workflow from our institute (https://github.com/ikmb/TOFU-MAaPO) that relies on BBtools and bowtie2 for mapping and host decontamination, and bioBakery tools (MetaPhlAn 4.0 and HUMaN 3.6) for taxonomic and functional potential profiling. Raw reads were quality trimmed and mapped to the mouse genome (GRCm39) to discern between sequenced reads from the host and the gut microbiome bacteria. Only samples containing >1 Gbp after trimming and host decontamination were kept in the analysis. Taxonomic features and microbial relative abundances were determined using MetaPhlAn 4.0 ^112^ with default parameters and its SGB database containing unique marker genes of 26,970 species-level genome bins (SGBs) derived from a collection curated 1.01M prokaryotic and metagenome-assembled genomes (MAGs). Functional profiles (stratified pathways, gene families and enzyme categories) were computed using HUMAnN 3.6 ^113^ with default parameters. All abundances were transformed to CPM (count per million) before any statistical analysis.

### Microbiome phylogenetic analyses of 16S and metagenomics data

Uni- and multivariate analyses were performed in R (v.4.2.1) under phyloseq ^111^ (v.1.40.0), vegan ^114^ (v.2.6-2) and MAasLin2 ^115^ (v.1.10.0). All samples for diversity analyses were normalized by rarefaction to the minimum shared read count to account for differential sequencing depth among samples. Relative abundance was calculated by dividing the number of reads for an ASV by the total number of sequences in the sample. Within sample diversity (alpha diversity) was explored using various diversity indexes (Chao1, Shannon, Simpson, Observed ASVs) on ASV abundance data coupled to significance testing (Wilcoxon signed rank test or Kruskal-Wallis tests). Between-sample diversity (beta diversity) as well as differences in diet groups and time were explored and visualized using a Principal Coordinate Analysis (PCoA) plot based on Bray-Curtis distances. Associations of microbiome composition to specified covariates were tested with the implementation of PERMANOVA models (using the adonis2 function from the vegan package). The P and R^2^ values were determined by 10,000 permutations using time. To explore variation between and within individuals per diet phase at the genus level, the intraclass correlation coefficient (ICC) was computed by the *ICCest* function of the ICC R package, using as input the longitudinal rarefied abundances, defining each individual mouse as a distinct class. ICC estimation uses the variance components from a one-way ANOVA (between-group variance and within-group variance). To detect differences in the abundance of microbial features (16S rRNA, metagenomics) or pathways (metagenomics) between the experimental groups over time, we built linear mixed models using the MaAslin2 ^116^ package that included time, diet group and individual mice as a random variable (*microbial abundances ∼ (1|mouse) + diet switch + time*). P values were corrected for multiple hypothesis testing using the Benjamin-Hochberg procedure, and a false discovery rate <0.05 was defined as the significant threshold. Features (taxa, or pathways) appearing in at least 20% of the samples were included in the analysis.

### Identification of genes involved in the bacterial transformation of Linoleic acid-derived oxylipins

Databases of known bacterial oxylipin transformation enzyme genes such as Epoxide hydrolases (∼66.000) and Lipooxigenases (∼7800) were generated using the NCBI protein database. All genes tagged with the following search criteria were included in the database. (i) Epoxide hydrolases (10^th^ of March, 2023): (epoxide hydrolase[Protein Name] OR (epoxide[All Fields] AND hydrolase[All Fields])) AND (bacteria[filter] AND refseq[filter]). (ii) Lipooxygenases (24^th^ of March, 2023): Lipoxygenase[All Fields] AND bacteria[filter]. The specific enzyme database (Epoxide hydrolases and Lipooxigenases) and the UniRef90 database ^117^ were input into the ShortBRED identify pipeline (v.0.9.5) and used to generate protein-specific markers ^118^. These markers along with the quality-controlled metagenomics samples were used as input for ShortBRED quantify to get the presence of copy numbers of the target protein markers.

### Non-targeted metabolomics using HILIC UHPLC-MS/MS

Fecal pellets (∼25mg) were weighed in sterile ceramic bead tubes (NucleoSpin® Bead Tubes, Macherey-Nagel, Dueren, Germany) and 1mL of methanol was added (−20°C; LiChrosolv, Supelco, Merck, Darmstadt, Germany). Metabolite extraction was performed with Precellys® Evolution Homogenizer with following settings (4,500 rpm, 40 seconds of 3 cycles with 2 seconds pause time, Bertin Corp., Rockville, Maryland, USA). Samples were centrifuged for 10 minutes at 21,000 × g, cooled at 4°C and 900 µL of supernatant was transferred into new tubes. Fifty microliters of supernatant were evaporated (40°C, SpeedVac Concentrator, Savant SPD121P, ThermoFisher Scientific, Waltham, MA, United States) and reconstituted in 75% acetonitrile (ACN; LiChrosolv, hypergrade for LC-MS; Merck KGaA). Samples were analyzed using an UHPLC system (ExionLC, Sciex LLC, Framingham, MA, USA) coupled to a quadrupole time-of-flight mass spectrometer (X500R QTOF MS, Sciex LLC, Framingham, MA, USA). Parameters of the MS for positive and negative electrospray ionization mode are summarized in Table S3 and hydrophilic interaction liquid chromatography (HILIC) was performed as described previously ^119^ to analyze small polar metabolites.

### Targeted metabolomics identification of 9-HODE and 12,13-diHOME

Identification and semi-targeted peak picking of 12,13-diHOME ((9Z,12S,13S)-12,13-dihydroxyoctadec-9-enoic acid) and 9-HODE was conducted with the same LC-MS method as described previously ^120^. Here, we used an UHPLC system (ExionLC, Sciex LLC, Framingham, MA, USA) coupled to a quadrupole time-of-flight mass spectrometer (X500R QTOF MS, Sciex LLC, Framingham, MA, USA). We analyzed 12,13-diHOME standard, present on the GUTMLS01-1EA plate (Sigma-Aldrich, Plate 1, Row A, final concentration of around 0.0025 µg/µL, dissolved and diluted in methanol) and fecal extracts of CONVR and GF mice. The identity of 9-HODE was verified by matching against spectral library of MassBank of North America. The X500R QTOF MS was set to negative electrospray ionization mode with parameters summarized in Table S3.

### Metabolomics data processing and analysis

Raw LC-MS (.wiff2) data were post-processed using GeneData Expressionist Refiner MS 15.0 (GeneData GmbH, Basel, Switzerland) including chemical noise subtraction (signal with intensities <200 were removed), chromatographic peak picking, chromatogram isotope clustering, valid feature filter (features present in 10% of samples and at least 500 maximum intensity), retention time range restriction (0.4-9 minutes), annotate known peaks (mass-to-charge tolerance of 0.005 Dalton and retention time tolerance of 0.1 minutes), MS/MS consolidation (keep signals with highest total ion count for one cluster in one sample), MS/MS peak detection and export to MASCOT generic files. Processing resulted in a data matrix containing clusters (*m*/*z* and retention time values) and observed maximum intensity values for each sample. Maximum intensity values were normalized to the wet fecal weight. For each cluster we kept MS/MS with highest total ion counts within one sample. MS/MS of clusters across samples were combined with the consensusSpectrum() function of MSnbase (), using mzd of 0.005, minProp of 0.1 and sum for intensityFun argument. Ions of MS/MS were matched with 0.005 Dalton, and intensities of aggregated peaks were summed, if ions were present in minimum 10% of matched spectra. Ions with intensities ≤500 and m/z values ≥ precursor m/z+2 were discarded from consensus spectra. Identification was done by matching experimental MS/MS spectra against spectral libraries, downloaded from MassBank of North America by using MS PepSearch (0.01 Dalton mass tolerance for precursor and fragment Search). Furthermore, GNPS molecular networking and GNPS Library Search was engaged for metabolite identification of experimental MS/MS. Within MS PepSearch output, matches with dot product over 500 were kept and multiple matches for one cluster were filtered for the best match. We kept clusters that are present in at least 4 samples and clusters that have valid MS/MS information or metabolite identities. With this approach, we kept 793 and 833 clusters for positive and negative mode, respectively. Furthermore, SIRIUS was used for metabolite identification and classification by an in-silico approach using MS/MS ^121^. Multivariate statistical analysis was then performed using Wilcoxon signed rank test implemented in R.

### Statistical analysis

Biostatistical analyses were performed using GraphPad Prism (version 8) software (GraphPad, Inc, La Jolla, CA) or R (v 3.2.5). Specific comparisons and analyses are described in the individual method sections. All pairwise comparisons were performed using Mann-Whitney U test and time-series multiple comparisons were done using Holm-Šídák’s multiple comparisons test. Data are shown as mean ± standard error of the mean (SEM).

## Declarations

## Acknowledgments

The authors thank Sabine Kock, Stefanie Baumgarten, Maren Reffelmann, Vivian Wegner, Tanja Klostermeier, Dorina Ölsner, Meike Hansen, Ronja Möhring and Sophie Reiher for excellent technical assistance. We further thank Hans-Christian Behrens for providing access to the Hamamatsu NanoZoomer S60 Digital Slide Scanner, Jacob Hamm for experimental support and Robert Häsler, Louise Thingholm, Eike Wacker and David Ellinghaus for valuable input regarding the RNAseq and metagenomic analyses. We thank Anthony Coll, John Tadross and Stephen Ó Rahilly, University Cambridge for insightful critical discussions and provision of reagents.

## Author contributions

FS, NM, NS, FS and PR designed research. FS, NM, VL, NS, JPB, AW, FH, TG, FG, LS, AB, SK, SW, GI, FT, FS and PR performed experiments and analyzed the data. SS, JMP, RSB, PSK, JFB provided critical resources. FS, NM, VL, JPB, FS and PR prepared the figures. FS and PR obtained funding. FS, NM, VL, JPB, FH, FS and PR co-wrote the manuscript with critical input from all authors. All authors read and approved the final manuscript.

---

**Figure S1:**
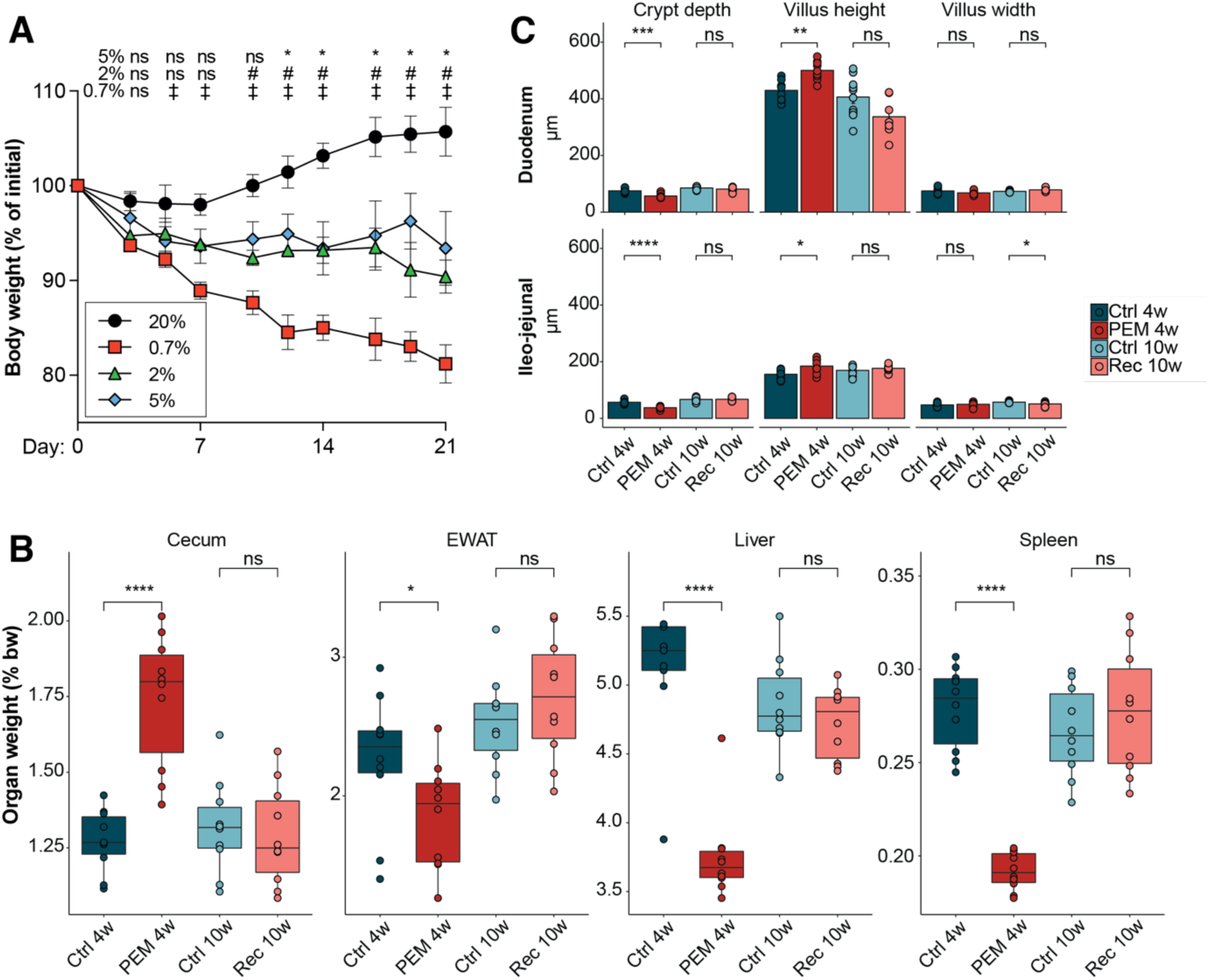
PEM drastically alters intestinal architecture and organ measures. **A)** Body weight development of adult mice fed either a control diet (20% protein) or PEM diets with various protein contents (0.7, 2 and 5%) for 3 weeks. **B)** Cecum, EWAT (epididymal white adipose tissue), liver and spleen weights relative to body weight of mice during the PEM diet and recovery intervention. **C)** Crypt depth, villus height and villus width from small intestine of mice during the PEM diet and recovery intervention.

**Figure S2:**
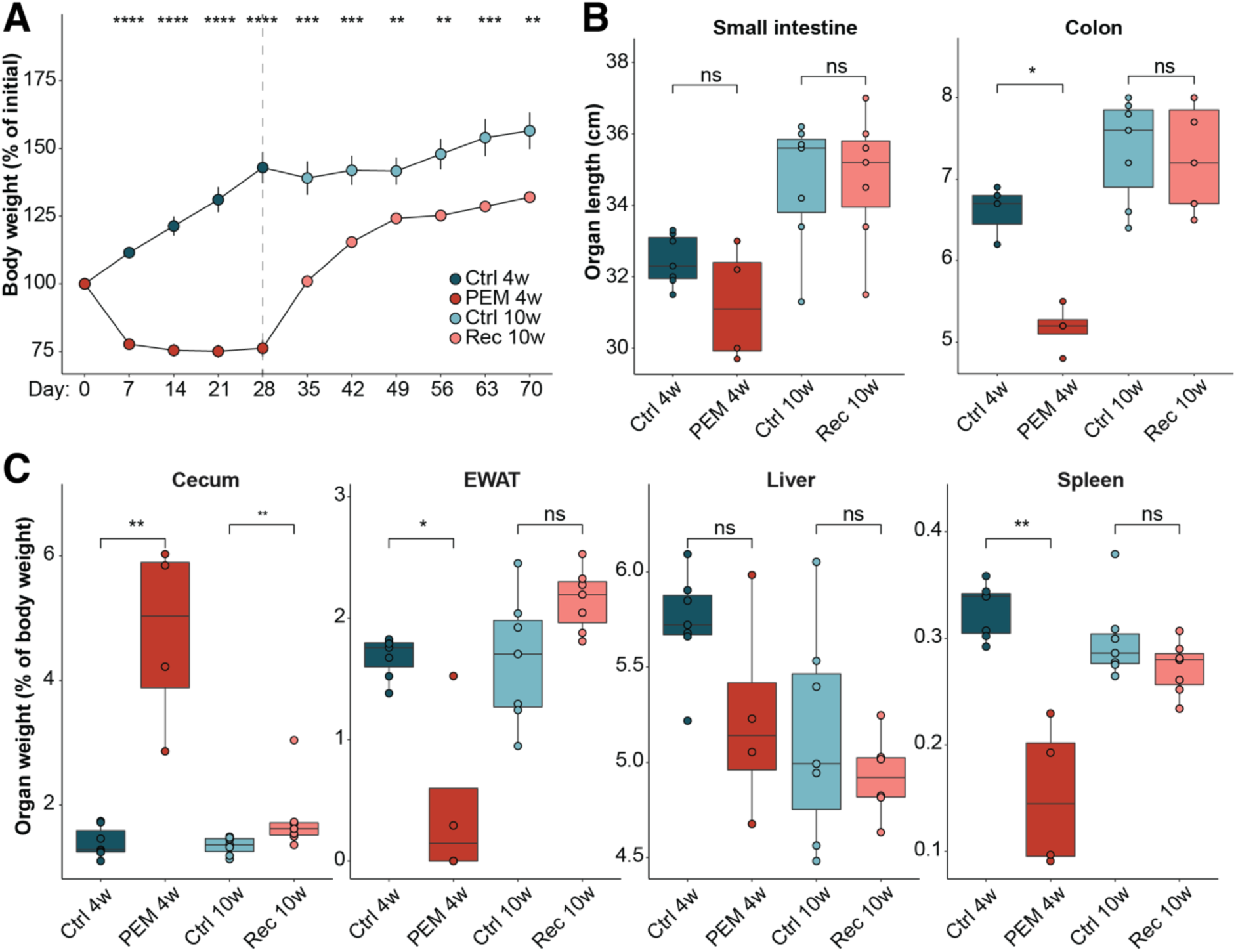
PEM during postnatal development aggravates the disease phenotype. **A)** Body weight curve of 4-week-old mice weaned onto PEM diet for 4 weeks and then switched to regular control diet for recovery. **p < 0.01, ***p < 0.001 and ****p < 0.0001 using two-way ANOVA. **B**) Lengths of the small and large intestine during PEM and recovery. *p < 0.05 and ns = not significant using Mann-Whitney U test. **C)** Cecum, EWAT (epididymal white adipose tissue), liver and spleen weights relative to body weight of mice during the PEM diet and recovery intervention. *p < 0.05, **p < 0.01 and ns = not significant using Mann-Whitney U test.

**Figure S3:**
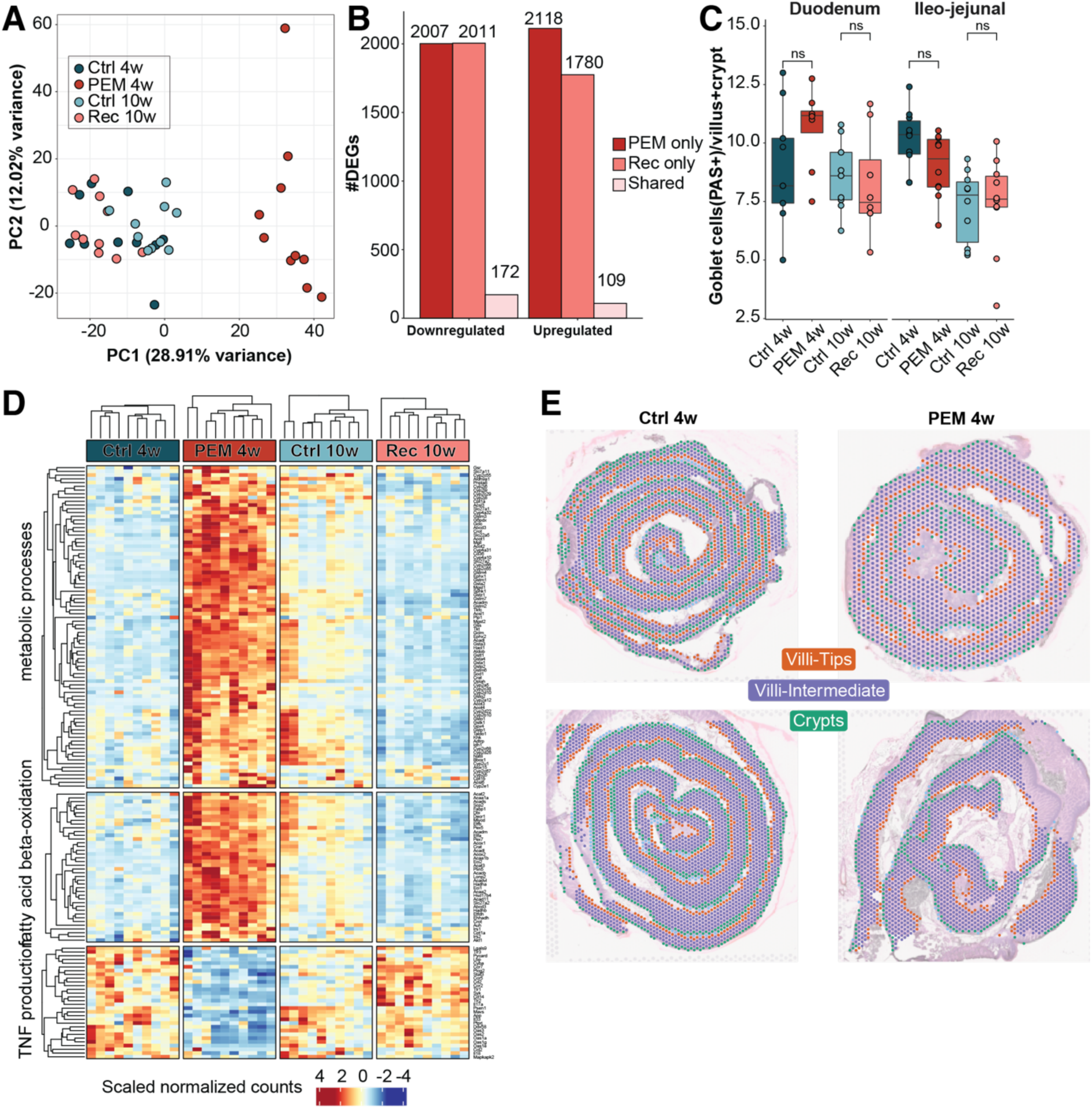
Differential regulation of genes in intestinal epithelial cells. **A)** Principal component analysis (PCA) for RNA sequencing samples. **B)** Number of significantly upregulated and downregulated genes in small intestinal epithelial cells from mice during acute PEM (4 weeks) and after recovery (10 weeks) and respective controls. **C)** Boxplots showing average number of goblet cells (PAS+) per villus and crypt in duodenum and ileo-jejunal regions of the small intestine of mice under Ctrl and PEM diets along with representative histological images. ns: *p* > 0.05 Ctrl_4w versus PEM_4w and Ctrl_10w versus Rec_10w using Mann-Whitney U test. **D)** Heatmap showing the expression of DEGs corresponding to selected GO terms enriched in upregulated and downregulated PEM only DEGs. Scaled normalized gene expression counts across all samples are plotted. **E)** Overlay of spatial transcriptomic spots classified by different regions on high-quality H&E stainings from PEM_4w and Ctrl_4w mice.

**Figure S4:**
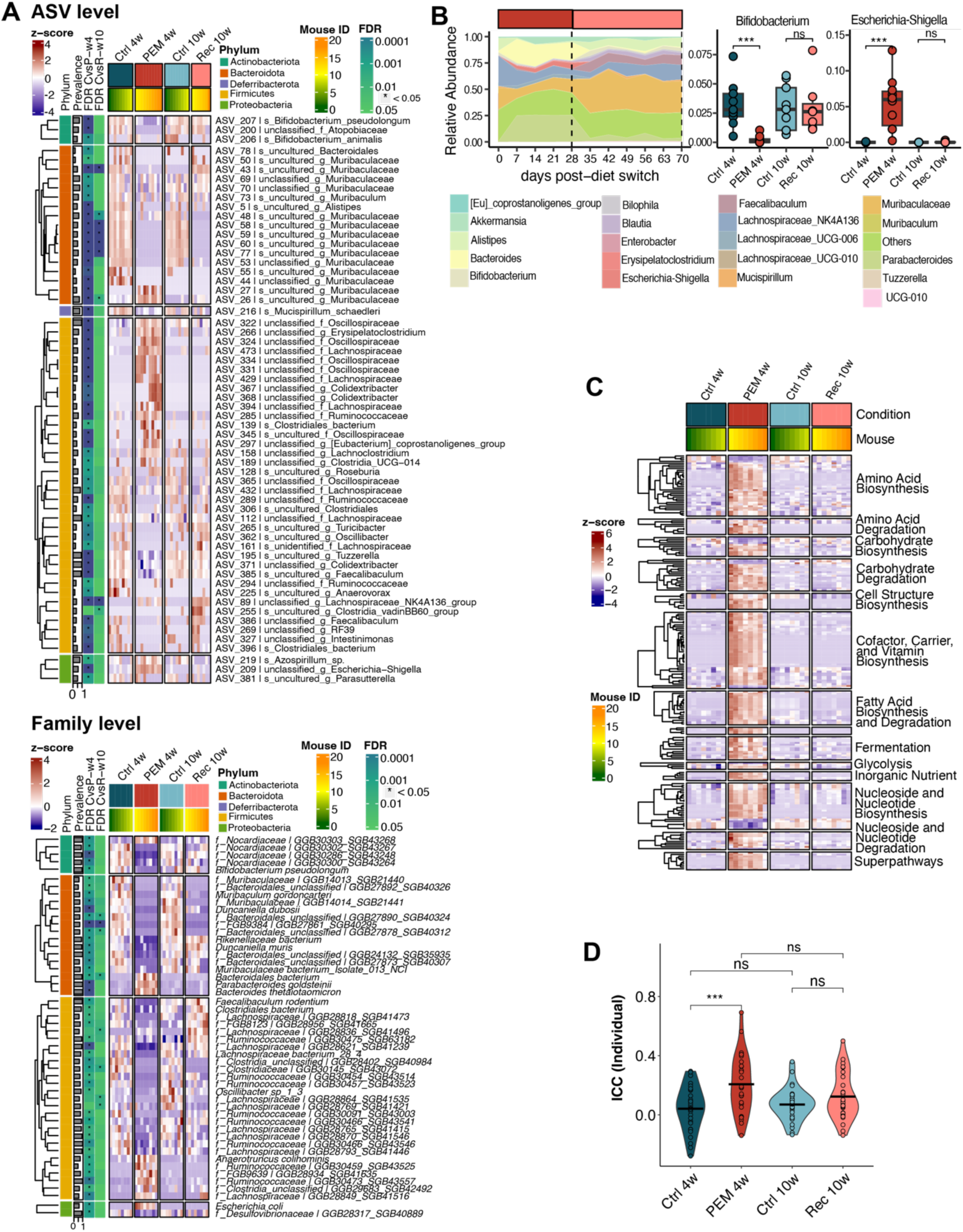
Longitudinal microbiome functions affected by PEM and recovery. **A)** Heatmap of relative abundance of bacterial species (ASV) and families with significant changes under acute PEM (PEM_4w vs Ctrl_4w) and after recovery (Rec_10w vs Ctrl_10w). For each cell, colors indicate the row-wise z-score of relative abundances, asterisks denote the FDR significance at each cross-sectional comparison and prevalence represents the percentage of non-zero features. Row-wise clusters represent features that belong to the same Phylum. **B)** Longitudinal area-stacked relative abundances of the top 20 significant genera identified by LefSe analysis. Unclassified genera and those with low relative abundance are grouped in “Others”. Relative abundances comparisons of two representative genera, *Bifidobacterium* and *Escherichia-Shigella*. Wilcoxon signed rank test was used to test significance. **C)** Heatmap of abundances of microbiome metabolic pathways (obtained from HUMAnN 3.6) with significant changes under acute PEM (PEM_4w vs Ctrl_4w) and after recovery (Rec_10w vs Ctrl_10w). For each cell, colors indicate the row-wise z-score of relative abundances, asterisks denote the FDR significance at each cross-sectional comparison and prevalence represents the percentage of non-zero features. Row-wise clusters represent features that belong to the same higher-level pathway. **D)** Intraclass correlation coefficient (ICC) of individual genera across diet phases. Statistical differences were assessed by Wilcoxon signed-rank test.

**Figure S5:**
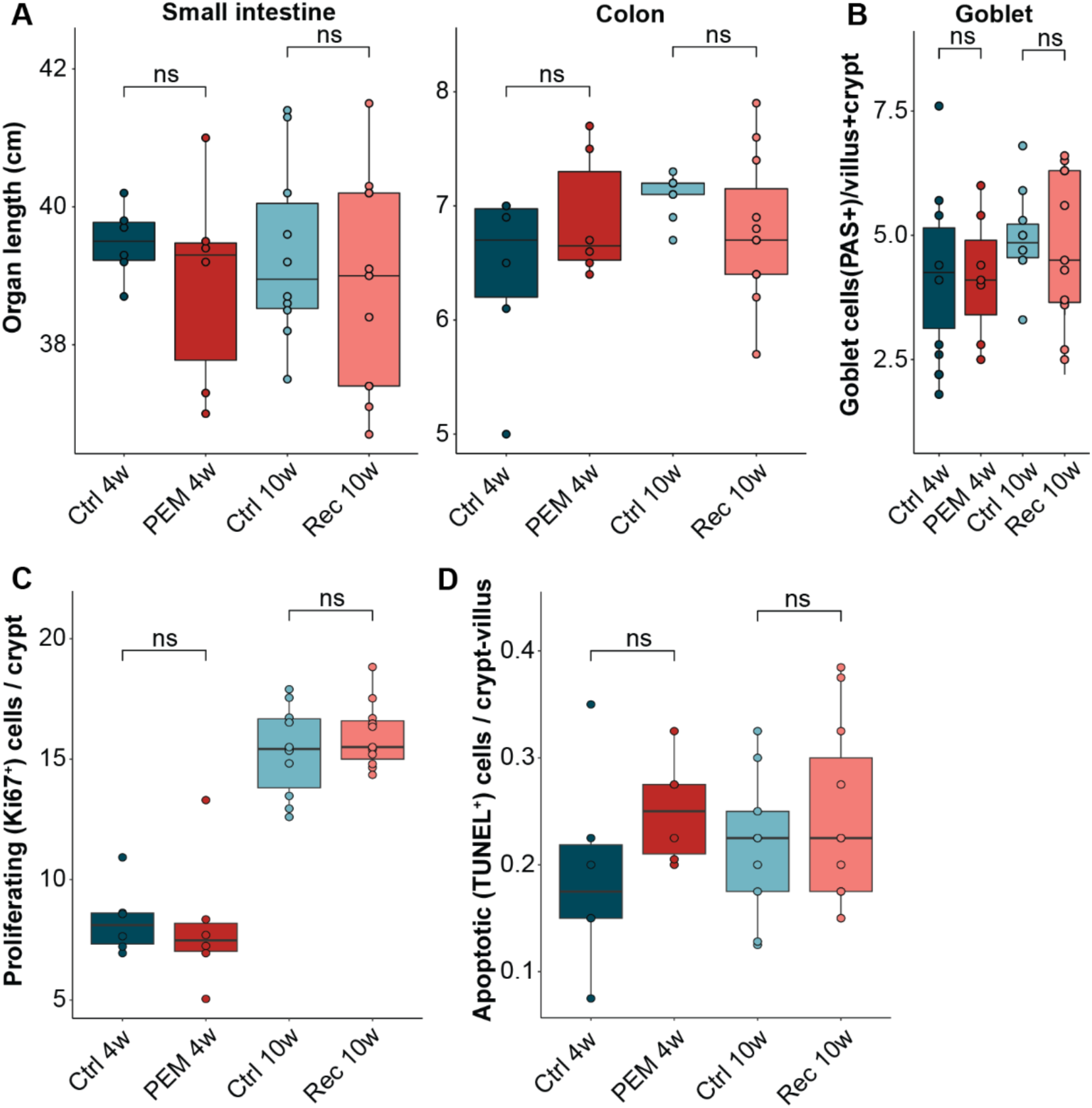
Ameliorated PEM phenotype in GF mice. GF mice (n=10-11 per group) were subjected to a dietary PEM-recovery intervention experiment and analyzed for organ measures and cell counts. **A)** Length of the small and large intestines. **B)** Number of PAS-positive goblet cells per villus-crypt-unit in intestinal small sections. **C)** Number of Ki67-positive proliferative epithelial cells per crypt. **D)** Number of TUNEL-positive apoptotic epithelial cells per crypt-villus unit. Significance testing was performed using Wilcoxon-Mann-Whitney-Test.

**Figure S6:**
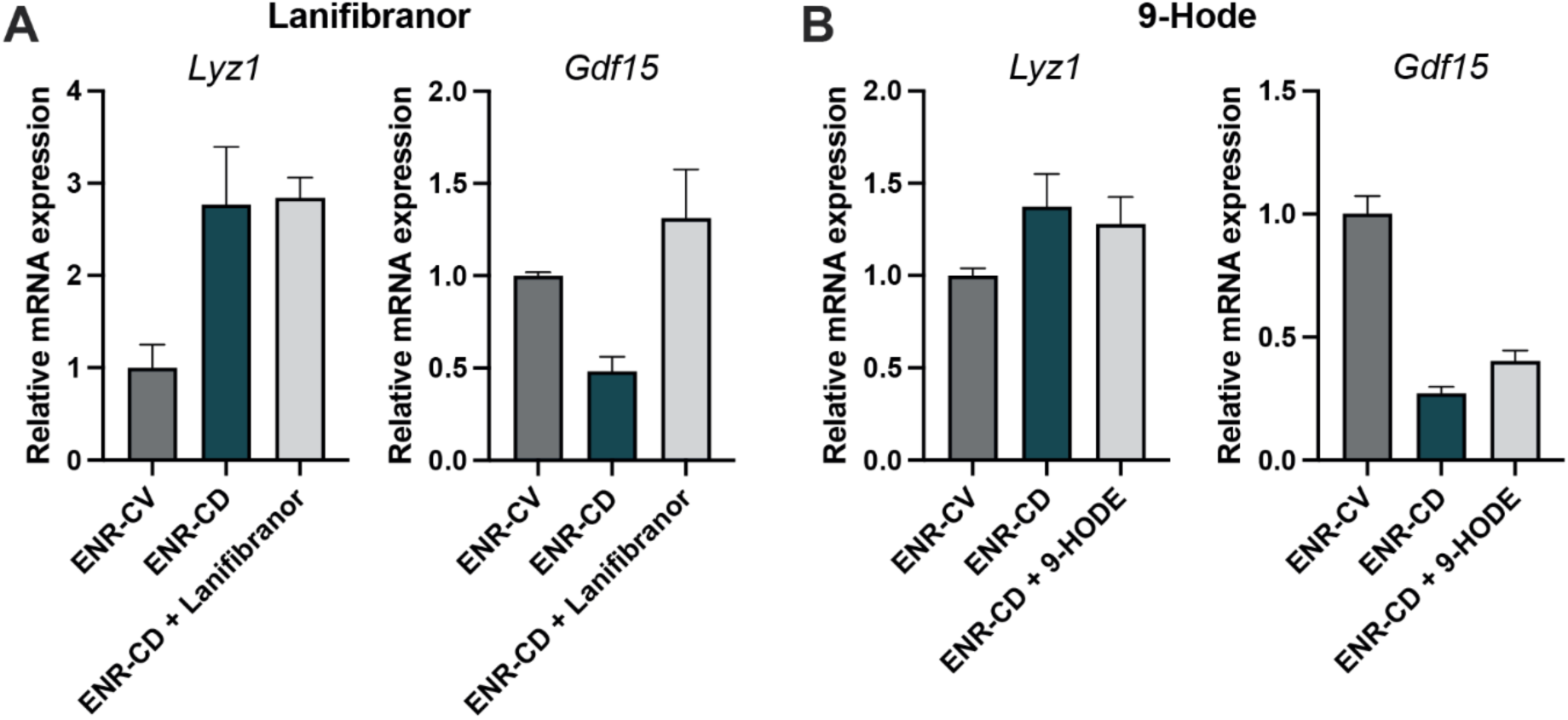
PPAR-activation or the microbial metabolite 9-HODE alone do with Microbiome status and PEM drive *Ppar* / *Gdf15* expression and Paneth cell differentiation. Expression of *Lyz1* and *Gdf15* were assessed in intestinal organoids stimulated with the **(A)** pan-PPAR-agonist Lanifibranor [20 µM] or the **(B)** lipid 9-HODE [1 µM] during Paneth cell differentiation. Note that stimulation with Lanifibranor or 9-HODE did not alter *Lyz1* or *Gdf15* expression in contrast to stimulation during PEM as shown in Figure 6.

**Table S1:** Formulation of PEM diets.

| **Ingredient** | **U** | **0.7 %**<br>**Protein** | **2.0 %**<br>**Protein** | **5.0 %**<br>**Protein** | **20.0 %**<br>**Protein** |
| --- | --- | --- | --- | --- | --- |
| Product No. (Ssniff) |  | *S0071-E710* | *S0071-E711* | *S0071-E712* | *S0071-E714* |
| **Casein** | **%** | **0.680** | **2.200** | **5.600** | **21.500** |
| L-Cystine | % | 0.100 | 0.100 | 0.100 | 0.300 |
| Maltodextrin | % | 32.020 | 30.500 | 27.100 | 11.000 |
| Corn starch, pre-gelatinized | % | 32.400 | 32.400 | 32.400 | 32.400 |
| Dextrose | % | 12.000 | 12.000 | 12.000 | 12.000 |
| Sucrose | % | 0.200 | 0.200 | 0.200 | 0.200 |
| Cellulose powder | % | 7.000 | 7.000 | 7.000 | 7.000 |
| Inulin powder | % | 2.300 | 2.300 | 2.300 | 2.300 |
| Vitamin premix | % | 1.000 | 1.000 | 1.000 | 1.000 |
| Mineral premix AIN | % | 5.000 | 5.000 | 5.000 | 5.000 |
| Choline Cl (50 %) | % | 0.250 | 0.250 | 0.250 | 0.250 |
| Dye [Y, G, B, V, R] | % | 0.050 | 0.050 | 0.050 | 0.050 |
| Soybean oil | % | 7.000 | 7.000 | 7.000 | 7.000 |
| ***Proximate contents*** |  |  |  |  |  |
| Crude protein | % | **0.7** | **2.0** | **5.0** | **19.0** |
| Crude fat | % | 7.0 | 7.0 | 7.0 | 7.0 |
| Crude fibre | % | 8.4 | 8.4 | 8.4 | 8.4 |
| Crude ash (minerals) | % | 4.2 | 4.2 | 4.2 | 4.3 |
| Carbohydrates | % | 76.5 | 75.1 | 71.7 | 56.1 |
| Energy (Atwater) | MJ/kg | 15.6 | 15.6 | 15.6 | 15.5 |
| kcal% Protein |  | 0.7 | 2 | 5 | 21 |
| kcal% Fat |  | 17 | 17 | 17 | 17 |
| kcal% Carbohydrates |  | 82 | 81 | 78 | 62 |

**Table S2:**
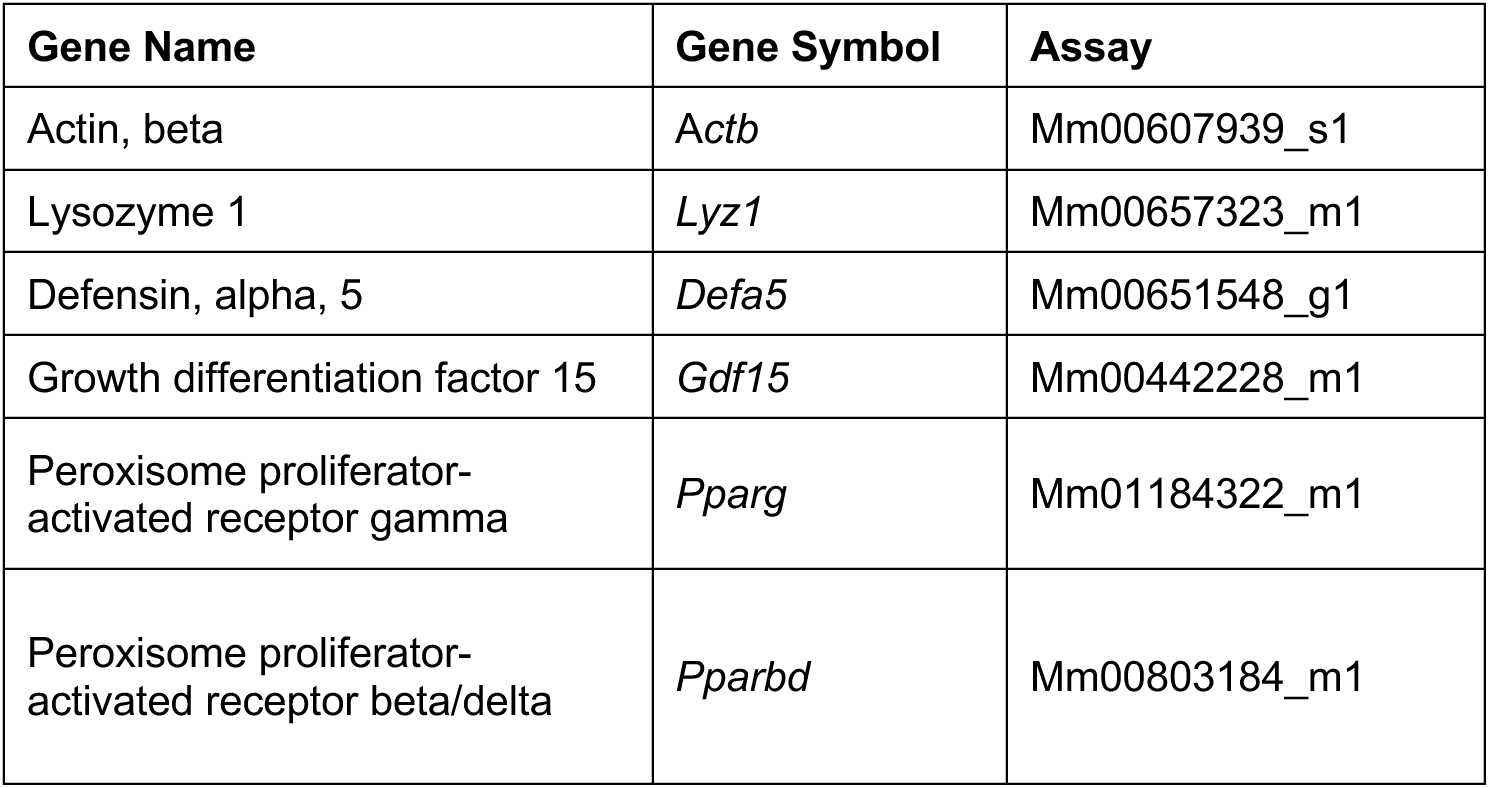
List of TaqMan^®^ Gene Expression Assays used in this study.

**Table S3:** X500R MS parameters for non-targeted metabolomics and targeted identification of fatty acids (12,13-diHOME and 9-HODE)

|  | Non-targeted metabolomics |  | Targeted identification | Unit |
| --- | --- | --- | --- | --- |
| Method duration (minutes) | 13.5 | 13.5 | 25 | min |
| Total Scan Time | 0.412 | 0.412 | 0.412 | s |
| Curtain gas | 30 | 30 | 30 | psi |
| Ion source gas 1 | 45 | 45 | 45 | psi |
| Temperature | 500 | 500 | 500 | °C |
| Ion source gas 2 | 45 | 45 | 45 | psi |
| CAD gas | 7 | 7 | 7 | - |
| Polarity | **Positive** | **Negative** | **Negative** | - |
| Spray voltage | 5500 | -4500 | -4500 | V |
| TOF start mass | 50 | 65 | 65 | Dalton |
| TOF stop mass | 1000 | 1000 | 1000 | Dalton |
| Declustering potential | 50 | -50 | -50 | V |
| DP spread | 0 | 0 | 0 | V |
| Accumulation time | 0.1 | 0.1 | 0.1 | s |
| Collision energy | 5 | -5 | -5 | V |
| CE spread | 0 | 0 | 0 | V |
| IDA Criteria | Small molecule | Small molecule | Small molecule | - |
| Maximum candidate ions | 10 | 10 | 10 | - |
| Intensity threshold | 1000 | 1000 | 1000 | cps |
| TOF start mass | 50 | 50 | 50 | Dalton |
| TOF stop mass | 1000 | 1000 | 1000 | Dalton |
| Declustering potential | 80 | -80 | -80 | V |
| DP spread | 0 | 0 | 0 | V |
| Accumulation time | 0.025 | 0.025 | 0.025 | s |
| Collision energy | 35 | -35 | -35 | V |
| CE spread | 15 | 15 | 15 | V |

